# *Candidatus* Nealsonbacteria (OD1) are biomass recycling ectosymbionts of methanogenic archaea in a stable benzene-degrading enrichment culture

**DOI:** 10.1101/2022.04.20.488981

**Authors:** Xu Chen, Olivia Molenda, Christopher T. Brown, Courtney R. A. Toth, Shen Guo, Fei Luo, Jane Howe, Camilla L. Nesbø, Christine He, Elizabeth A. Montabana, Jamie H. D. Cate, Jillian F. Banfield, Elizabeth A. Edwards

## Abstract

The Candidate Phyla Radiation (CPR) is a very large group of bacteria with no pure culture representatives, first discovered by metagenomic analyses. Within the CPR, candidate phylum Parcubacteria (previously referred to as OD1) within the candidate superphylum Patescibacteria is prevalent in anoxic sediments and groundwater. Previously, we had identified a specific member of the Parcubacteria (referred to as DGGOD1a) as an important member of a methanogenic benzene-degrading consortium. Phylogenetic analyses herein place DGGOD1a within the *Candidate* clade Nealsonbacteria. Because of its persistence over many years, we hypothesized that *Ca*. Nealsonbacteria DGGOD1a must serve an important role in sustaining anaerobic benzene metabolism in the consortium. To try to identify its growth substrate, we amended the culture with a variety of defined compounds (pyruvate, acetate, hydrogen, DNA, phospholipid), as well as crude culture lysate and three subfractions thereof. We observed the greatest (10 fold) increase in the absolute abundance of *Ca*. Nealsonbacteria DGGOD1a only when the consortium was amended with crude cell lysate. These results implicate *Ca*. Nealsonbacteria in biomass recycling. Fluorescent in situ hybridization and cryogenic transmission electron microscope images revealed that *Ca*. Nealsonbacteria DGGOD1a cells were attached to larger archaeal *Methanothrix* cells. This apparent epibiont lifestyle was supported by metabolic predictions from a manually curated complete genome. This is one of the first examples of bacterial-archaeal episymbiosis and may be a feature of other *Ca*. Nealsonbacteria found in anoxic environments.

## Introduction

The candidate phylum Parcubacteria belongs to a superphylum referred to as the Candidate Phyla Radiation (CPR) which was first defined by culture-independent metagenomic analysis (1). Members of the CPR are phylogenetically diverse and ubiquitous, but are especially abundant in lakes, sediment, and groundwater (1, 2). Although their roles are still not well characterized, clues are emerging from the rapidly growing number of metagenome sequencing projects and genome reconstructions. To date, most members of the Parcubacteria remain uncultured and are only known from metagenome assembled genomes (MAGs). In 2020, Chen *et al*. (3) summarized 38 closed, complete MAGs or cMAGs belonging to the CPR. As of 2022, 19 cMAGs from the Parcubacteria within the CPR can be retrieved from NCBI, but none is for a member of the Nealsonbacteria subgroup. The genomes of Parcubacteria are extremely small (~1 Mb) and include many hypothetical proteins which lack homologs in well-studied microbes (4). The reduced genomes of Parcubacteria encode few metabolic capacities, lacking the ability to synthesize lipids and often missing complete pathways for the biosynthesis of several vitamins, amino acids, and nucleotides (5, 6). Coupled with a lack of an electron transport system, the small cells typical of Parcubacteria and other members of the CPR are thought to obtain nutrients and energy through close contact with an obligate host or partner cell (7). Several recent publications support this hypothesis. For example, Huddy et al. (8) inferred an association between CPR member Saccharibacteria (TM7) and Actinobacteria based on abundance patterns. He at el. (9) reveled episymbiotic interaction between ultrasmall cells and hosts in groundwater by Cryo-TEM. Scanning transmission x-ray microscopy (STXM) data obtained by Alvarado et al. (10) showed an association of ultra-small cells with filamentous bacteria which encapsulated with elemental sulfur spherical granules. The most definitive study was a study of Saccharibacteria (TM7) and its bacterial host, *Actinomyces odontolyticus* XH001, in cultivated co-cultures (11). Saccharibacteria genomes share similarities with the Parcubacteria in that they are small and lack complete biosynthetic pathways. Metagenomic analyses suggest that Parcubacteria may also be closely associated with a host microbe, perhaps as an episymbiont (5, 9, 12, 13) but experimental evidence is scant because of the difficulty in cultivating and enriching these tiny microorganisms from complex ecosystems.

Interestingly, a stable methanogenic benzene-degrading enrichment culture maintained in our laboratory, referred to as the OR consortium (for Oil Refinery), provides an opportunity to study a member of the Parcubacteria in vivo. The OR consortium was originally derived from oil-contaminated sediments in 1995 and has since been maintained in anaerobic minimal medium and provided with benzene as the only added electron donor and carbon source (14). Multiple subcultures have been established over the years, including a scaled-up lineage currently being used for bioaugmentation field trials (15–17). The dominant microbes in these subcultures include specialized benzene-fermenting *Desulfobacterota* (ORM2), several methanogenic archaea, and - most interestingly for this study - a member of the Parcubacteria originally referred to as OD1 but now further classified as *Candidatus* Nealsonbacteria (Fig. S1) in this study. The consortium repeatedly degrades benzene to methane and CO_2_ at rates from 0.2 to 10 mg benzene/L/day, however the biochemical mechanism for anaerobic benzene biodegradation is still poorly understood. Toth et al. (17) recently described the current state of knowledge of the OR consortium, highlighting the coordinated metabolic interactions of ORM2 and acetoclastic and hydrogenotrophic methanogens for complete mineralization of benzene. *Ca*. Nealsonbacteria might also play role in the OR consortium because they have consistently been detected in every culture survey (18) although a clear correlation between the abundance of *Ca*. Nealsonbacteria and benzene degradation rate has not been found. OR consortium cultures are maintained in fed-batch mode with infrequent (every 1-2 months) shallow transfers into fresh medium resulting in long cell residence times. Despite years of maintenance solely on benzene the total microbial concentration remains below 10^9^ cells/mL (19). Accumulated dead cells are a likely an alternative carbon and energy source for microbial growth. More generally, *Ca*. Nealsonbacteria are also found in anoxic hydrocarbon- and organohalide-contaminated sites (20, 21), high pH spring water (22), and other nutrient-limited groundwater environments (9), where biomass turnover rates are sometimes high, perhaps in response to stress, therefore we hypothesized that these microbes may play a role in biomass recycling.

We sought to identify the substrates of *Ca*. Nealsonbacteria and any possible host associations within the benzene-degrading OR consortium using enrichment experiments on various substrates, quantitative PCR (qPCR), sequencing and genome reconstruction, and various types of microscopy. The most growth of *Ca*. Nealsonbacteria was seen in cultures amended with crude cell lysate (prepared from the consortium itself or from *Escherichia coli*). Fluorescence *in situ* hybridization (FISH) and electron microscopy (EM) revealed *Ca*. Nealsonbacteria as an epibiont on the surface of *Methanothrix*, a methanogenic archaeon essential for completing the biotransformation of benzene to methane. Hybrid assemblies of short read (Illumina) and long read (PacBio) metagenomic sequences were used to successfully reconstruct the complete closed genome of the uncultured Parcubacteria in OR consortium, designated DGGOD1a. This 1.16 Mbp genome displays biosynthetic deficiencies and other episymbiotic traits as observed in most other Parcubacteria genomes. A maximum likelihood concatenated ribosomal protein tree placed the DGGOD1a genome within the candidate group Nealsonbacteria (Figure S1) as defined in Anantharaman et al. (23). These observations begin to shed light on uncharacterized members of the CPR and bring into focus the phenomenon of bacterial episymbiosis on archaeal hosts.

## Results and Discussion

### qPCR reveals abundance of *Ca*. Nealsonbacteria in methanogenic cultures

A qPCR survey of 15 different methanogenic benzene-degrading subculture bottles was conducted to measure the abundance of benzene-fermenting *Desulfobacterota* ORM2 and *Ca*. Nealsonbacteria (referred to as OD1 in previous studies), as well as the abundance of total bacteria and total archaea. One subculture bottle named “OR-1bBig” was selected for use in two substrate “Donor Trial” experiments because it contained a relatively high abundance of *Ca*. Nealsonbacteria (~5% based on 16S rRNA amplicon sequencing results) and sufficient culture volume was available (1.5 L). Other than benzene, the products of cell decay could feasibly serve as electron donors in these bottles. This led us to hypothesize that components of lysed cells might serve as growth substrates for *Ca*. Nealsonbacteria.

### *Ca*. Nealsonbacteria is most enriched in cultures amended with lysed cells

In Donor Trial #1, we tested three common substrates (pyruvate, acetate, and hydrogen), as well as French-Press lysed benzene culture, and two specific components of cells (salmon sperm DNA, and the phospholipid L-α-phosphatidylethanolamine) to see if any of these putative electron donors could support the growth of *Ca*. Nealsonbacteria, directly or indirectly (Table S1). We found that *Ca*. Nealsonbacteria concentrations increased in vials amended with lysed culture (Figure 1a). In these vials, *Ca*. Nealsonbacteria relative abundance increased from 5% to 30% and absolute abundance by an order of magnitude to 5.1×10^6^ copies (cells)/mL over 4 months (Figure 1; data in Table S2a). None of the other electron donors tested resulted in discernable enrichment of *Ca*. Nealsonbacteria relative to benzene-amended controls. In bottles amended with pyruvate, *Ca*. Nealsonbacteria absolute abundance increased ~5 times (up to 9.8×10^5^ copies/mL), but relative abundance decreased to 1% because other microbes grew much more (Figure 1b; Table S3). *Ca*. Nealsonbacteria growth in pyruvate and acetate amended bottles may have resulted from the direct substrate utilization or as an indirect consequence of cell growth and corresponding increase in dead cells. Either way, neither pyruvate nor acetate were selective for *Ca*. Nealsonbacteria. As expected, the abundance of benzene-degrading Desulfobacterota ORM2 dropped significantly in the absence of benzene (Figure 1b). In the vials amended with cell lysate (second bar; Figure 1b), in addition to enrichment of *Ca*. Nealsonbacteria to 30% of the population, we also observed enrichment of *Syntrophaceae* while the absolute abundance of methanogenic archaea was relatively stable (Table S2a). The ASV corresponding to *Syntrophaceae* was found to share 100% 16S RNA gene sequence identity to an uncultured *Syntrophaceae* (MH665869) from a methanogenic pyrite forming culture fed with ferrous sulfide (FeS), H_2_S, and CO_2_ (24). A similar metabolism may be occurring as our medium is reduced with FeS and buffered with bicarbonate (25).

**Figure 1.**
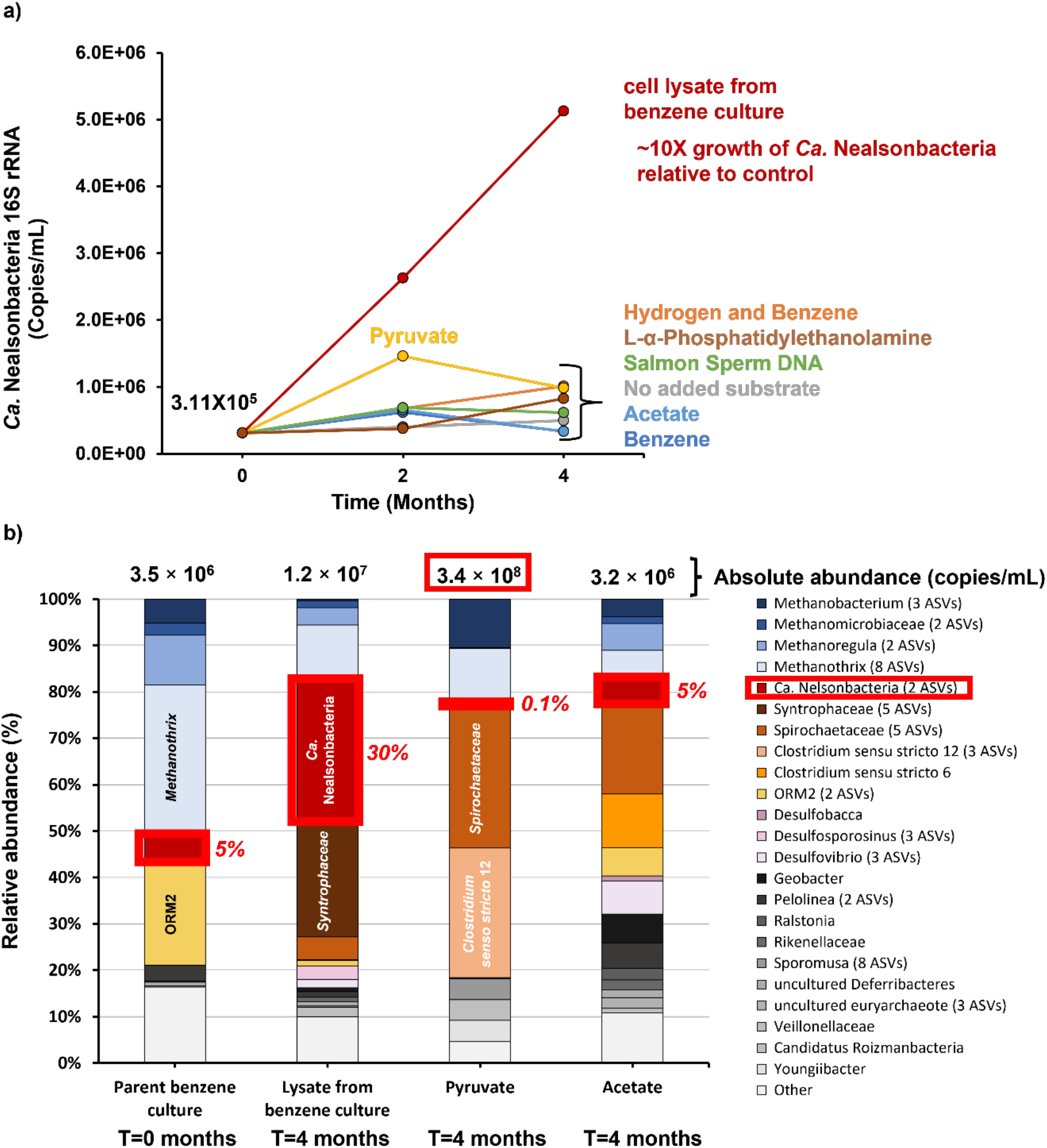
Results of Growth Trial #1. a) Absolute *Ca*. Nealsonbacteria abundance measured by qPCR in copies/mL, and b) microbial community profiles (relative abundance) as a function of substrate after 4 months of incubation, with total abundance shown above bars. In panel a) one DNA sample was collected on day 1 to represent initial conditions. Subsequent DNA samples were collected by sacrificing one entire bottle at each time point for each treatment due to the limited culture volume. In panel b) ASVs with relative abundances < 1% were combined as “Other”. ASVs with the same taxonomy at the genus level were combined. The number of combined ASVs for each taxon is shown in brackets. Absolute abundance was obtained by adding the qPCR results for total bacteria, total archaea and *Ca*. Nealsonbacteria together. “Parent Benzene culture” column is a 10% transfer of the parent culture at time zero.

Nonmetric dimensional scaling (NMDS) was applied to further analyze Donor Trial #1 ASV data (Figure S2a). The unfed “starved control” culture community clustered with communities from culture vials amended with cell lysate, consistent with products of cell decay supporting the microbial community in both cases. Methane production data from each treatment indicated that the added substrate equivalents (chemical oxygen demand [COD]) were consumed by 4 months (Table S4). Therefore, data collected at 7 months were from cultures that were starved for more than 3 months and marked with “+” in the NMDS analysis. Interestingly, we found that after 7 months, the communities did not converge. We expected to see enrichment of *Ca*. Nealsonbacteria in every vial after prolonged starvation. However, such enrichment was only clearly observed in acetate and pyruvate-amended vials (Figure 2). Perhaps there was insufficient biomass growth, therefore insufficient dead biomass in the other vials to detect enrichment of *Ca*. Nealsonbacteria. Indeed, total cell concentrations were highest in acetate and sodium pyruvate amended vials. Interestingly, a second ASV belonging to *Ca*. Nealsonbacteria (designated ASVb) was enriched in the acetate- and pyruvate-amended cultures (Figure 2) and became more abundant than *Ca*. Nealsonbacteria ASVa – the strain that is typically found in the benzene-degrading consortium (18). These two *Ca*. Nealsonbacteria ASVs share only 91.2% 16S rRNA sequence identity. This intriguing result suggests that different species may exhibit distinct preferences for the type of lysed cell material they can metabolize, because the acetate- and pyruvate-amended cultures selected for very different communities.

**Figure 2.**
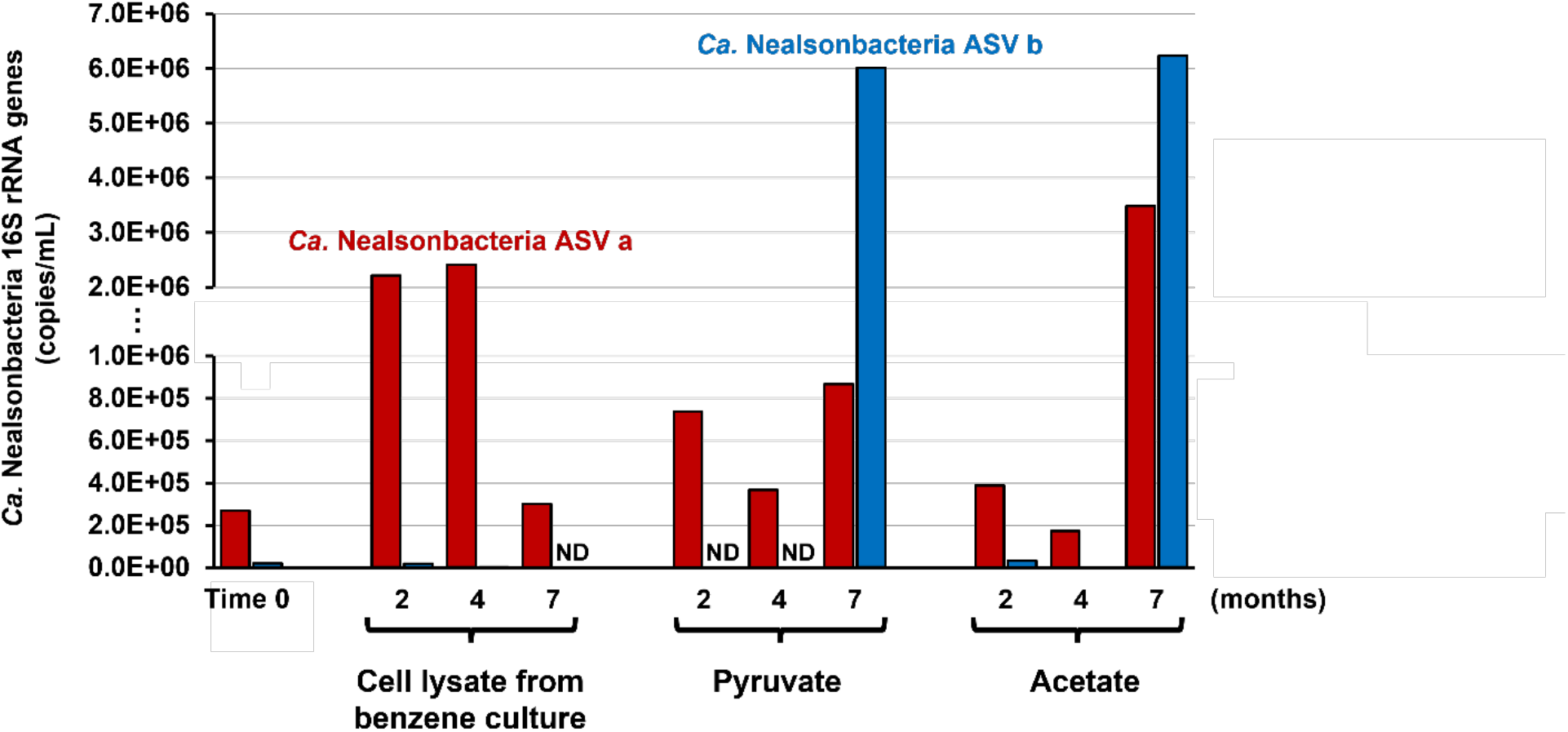
*Ca*. Nealsonbacteria strain abundance over 7 months of incubation. Figure shows treatments amended with benzene culture lysate, acetate, or pyruvate from Donor Trial #1. DNA samples were collected by sacrificing one entire bottle at each time point for each treatment due to the limited culture volume. Absolute abundance of *Ca*. Nealsonbacteria strains ASVa and ASVb were calculated by multiplying the relative abundance of these ASVs (obtained from amplicon sequencing) by the absolute abundance of total bacterial.

Building on the first trial, a second trial (Donor Trial #2) compared the growth of *Ca*. Nealsonbacteria on lysates from the benzene culture as well as *E. coli* lysate. Further, we tested three sub-fractions of the benzene culture lysate, which were separated by centrifugation and tangential flow filtration with a 50kDa cut off as depicted in Figure S3. The pyruvate treatment from Donor Trial #1 was also repeated but this time with added antibiotics (1 mM each kanamycin and vancomycin) to try to increase selectivity for *Ca*. Nealsonbacteria. The results of Donor Trial #2 (raw data in Table S2b) were similar to those of Donor Trial #1, with whole cell lysate resulting in significant (>10×) growth (Figure S4) regardless of source (*E. coli* or benzene culture). At first glance, the *E. coli* crude lysate appeared to be most stimulatory, but we later realized it was provided at 10× the concentration of the lysate from the benzene culture based on measured COD (Table S1c). The fact that *E. coli* lysate also supported the growth of *Ca*. Nealsonbacteria ruled out involvement of culture specific substrates (e.g., archaeal lipids). The sub-fractions of benzene culture lysate also resulted in enrichment although we observed considerable variability between duplicates (Figure S4). Of the three cell lysate fractions, the >50 kDa fraction (supernatant retentate) was most effective at enriching *Ca*. Nealsonbacteria after 4 months of incubation which may suggest that larger molecules may be important for growth. An NMDS analysis was also conducted on amplicon sequencing data from Donor Trial #2 (Figure S2b). These results show that the microbial communities enriched on each fraction of benzene culture lysate were slightly different after the first two months (triangles), but that they converged after four months (squares). Perhaps after four months, only larger, more slowly degraded biomolecules remained in all fractions and results in similar communities.

To look more closely at the microbial composition in each treatment, a heatmap of absolute abundance was generated showing the top 18 ASVs in Donor Trial #2 (Figure 3); raw data are reported in Table S5. *Ca*. Nealsonbacteria ASVa was the most abundant sequence variant detected. Only two other microbes (*Methanothrix* and *Spirochaetaceae*) were consistently enriched with *Ca*. Nealsonbacteria in vials amended with cell lysate, regardless of source or sub-fraction tested. Vials amended with lysed *E. coli* also enriched for ASVs belonging to *Clostridium* and *Bacteroides* that were not enriched in other treatments (Figure 3). Comparing the culture amended with only pyruvate in Donor Trial #1, the vials amended with antibiotics kanamycin and vancomycin successfully inhibited the growth of most microbes including *Ca*. Nealsonbacteria and *Methanothrix*. This indicates that *Ca*. Nealsonbacteria is antibiotic-sensitive or indirectly affected by its antibiotic-sensitive metabolic partners. It is interesting to note that the culture’s benzene fermenter (ORM2) seems to have persisted in those treatments. The growth of *Methanomethylovorans* and *Methanobacterium* in the presence of antibiotics confirms their resistance and together these data point to a possible means of enriching for benzene-degrading *Desulfobacterota* ORM2.

**Figure 3.**
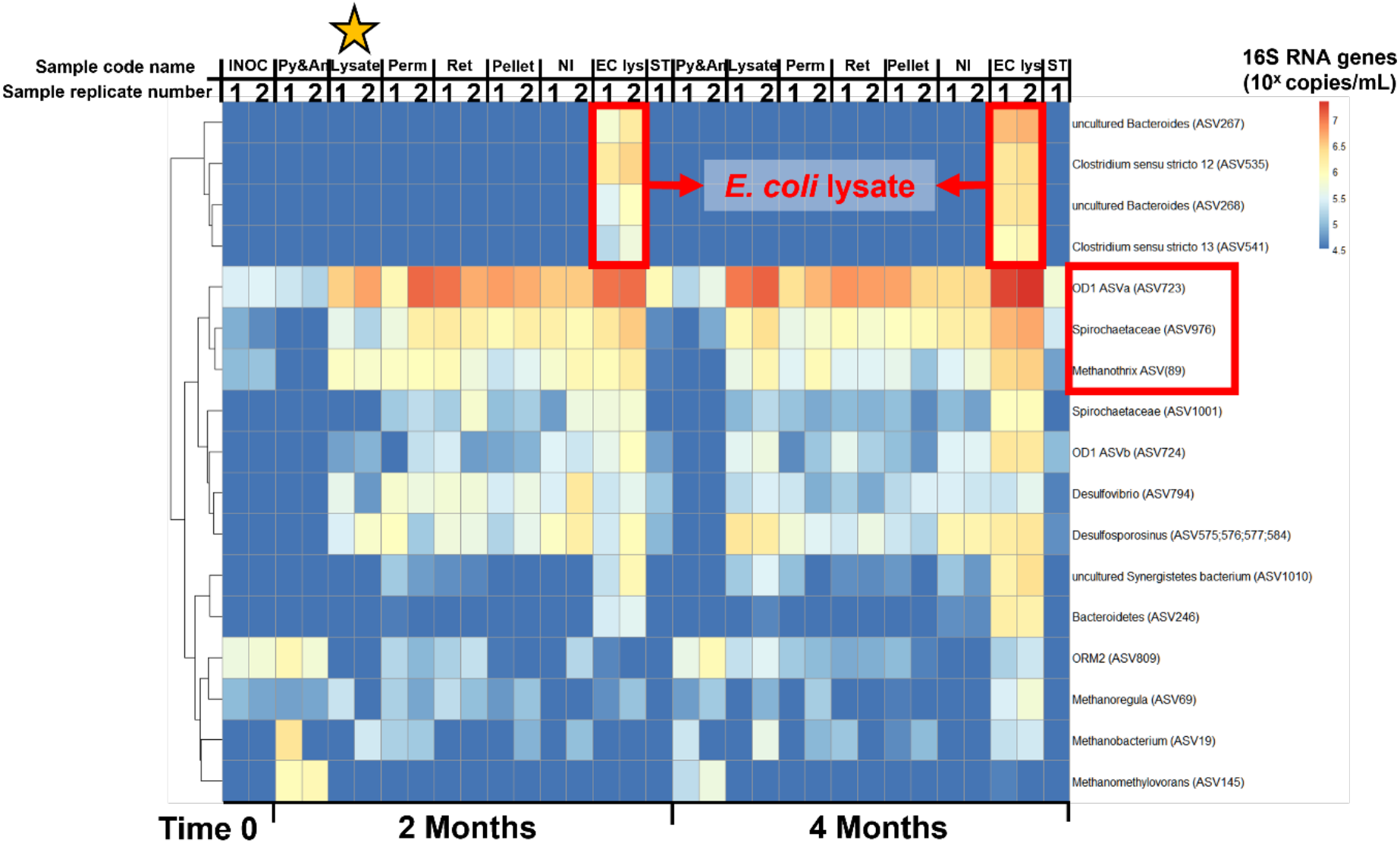
Heatmap of absolute abundance of top 20 ASVs in growth Trial #2. Top 20 ASVs were selected based on the sum of absolute abundance across all DNA samples collected. The ASVs under *Desulfosporosinus* are combined. Rows were clustered using a hierarchical clustering algorithm “hclust” in R based on similarity of absolute abundance. The columns represent different treatment bottles: INOC: Bottle with only 10% inoculum at time 0; NI: no inoculum (lysate only); Py&An: pyruvate & antibiotics; Lysate: French pressed benzene culture; Perm: lysate supernatant permeate through 50 kDa membrane; Ret: lysate supernatant retentate; Pellet: lysate pellet after centrifugation at 13,000 x g for 20 min; EC lys: French pressed lysate of *E. coli* culture; ST: Starved culture. The bracketed ASV numbers in the row names represent ordinal ASV number (see Table S3).

The two donor trials confirmed that compound(s) present in bacterial cell lysate can enrich *Ca*. Nealsonbacteria. The exact identity of the compound(s) remains elusive, but the data point to higher molecular weight and/or more recalcitrant proteins or extracellular polymeric substances, as these would present in all of the fractions tested. Two very distinct strains of *Ca*. Nealsonbacteria grew in the culture depending on treatment revealing subtle ecological preferences yet to be discovered.

### Fluorescence in situ microscopy (FISH) reveals *Ca*. Nealsonbacteria DGGOD1a as an epibiont of *Methanothrix*

To visualize the enriched *Ca*. Nealsonbacteria in the methanogenic benzene degrading culture by microscopy, we attempted to develop specific mono-fluorescent *in situ* hybridization (FISH) probes targeting various single stranded regions in the 16S rRNA gene. However, none of the probes worked, possibly because of low ribosome counts in these cells (26) or non-ideal targeting in the secondary structure. Very recently, Kuroda et al. (27) successfully designed and validated a mono-FISH probe for a *Ca*. Nealsonbacteria with a nearly identical 16S RNA gene sequence by targeting a different location on the gene; clearly this new FISH probe will be tested in the future.

As an alternative to a *Ca*. Nealsonbacteria-specific FISH probe, we used a combination of a universal Archaea FISH probe (ARCH915, Table S6) and DAPI staining to distinguish methanogens from bacteria. Given that *Ca*. Nealsonbacteria DGGOD1a was the most abundant bacterial phylotype in several of our lysate-amended enrichment cultures (Figure 3), we compared microscopic cell counts to qPCR abundance estimates of the same sample to confirm that the ultra-small cells were *Ca*. Nealsonbacteria. We focused primarily on one specific cell lysate enrichment culture (Lysate replicate #2 at 2 months) because of its high *Ca*. Nealsonbacteria abundance (>10^6^ cells/mL in this culture compared to all other bacterial phylotypes (Figure 3). In this *culture*, we observed many DAPI-stained ultra-small (~0.2 μm) cells attached to a single filamentous structure that stained with the archaeal FISH probe (Figure 4a). The ultra-small cell size matches the typical size of CPR members (5, 9) and is presumed to be *Ca*. Nealsonbacteria. The shape of the filamentous archaea is distinctive of *methanothrix* cells. We counted the number of small cells attached to long archaeal filaments as well as the number of filamentous archaea. We also counted the number of free-living microbes with similar size and shape to that of the attached small cells. Twenty-one fields of view were imaged and counted for each of two independent samples from a *Ca*. Nealsonbacteria enrichment culture. These small cells were by far the most numerous bacteria in all views, and thus must correspond to *Ca*. Nealsonbacteria, as it was most numerous via qPCR in this culture. The count of small *Ca*. Nealsonbacteria cells was 3.2-6.6×10^6^ cells /ml with attached cells ~2.5 times more abundant than free-living cells. (Table S7). Gene copies by qPCR in samples of DNA extracted from the same culture using primers specific for *Ca*. Nealsonbacteria gave a comparable result of 8.3×10^6^ copies/ml. Thus, we are confident that the small cells attached to the archaeal filaments are *Ca*. Nealsonbacteria. *Methanothrix* cell counts were also estimated at 1.9-3.4×10^6^ cells/mL, which also matches reasonably well to the value of 0.9 ×10^6^ cells/mL from qPCR results (Table S7). Interestingly, a similar association of *Ca*. Patescibacteria and *Methanothrix* was very recently reported in an artificial mixture of a culture of *Methanothrix soehngenii* GP6 with wastewater biosludge (27).

**Figure 4.**
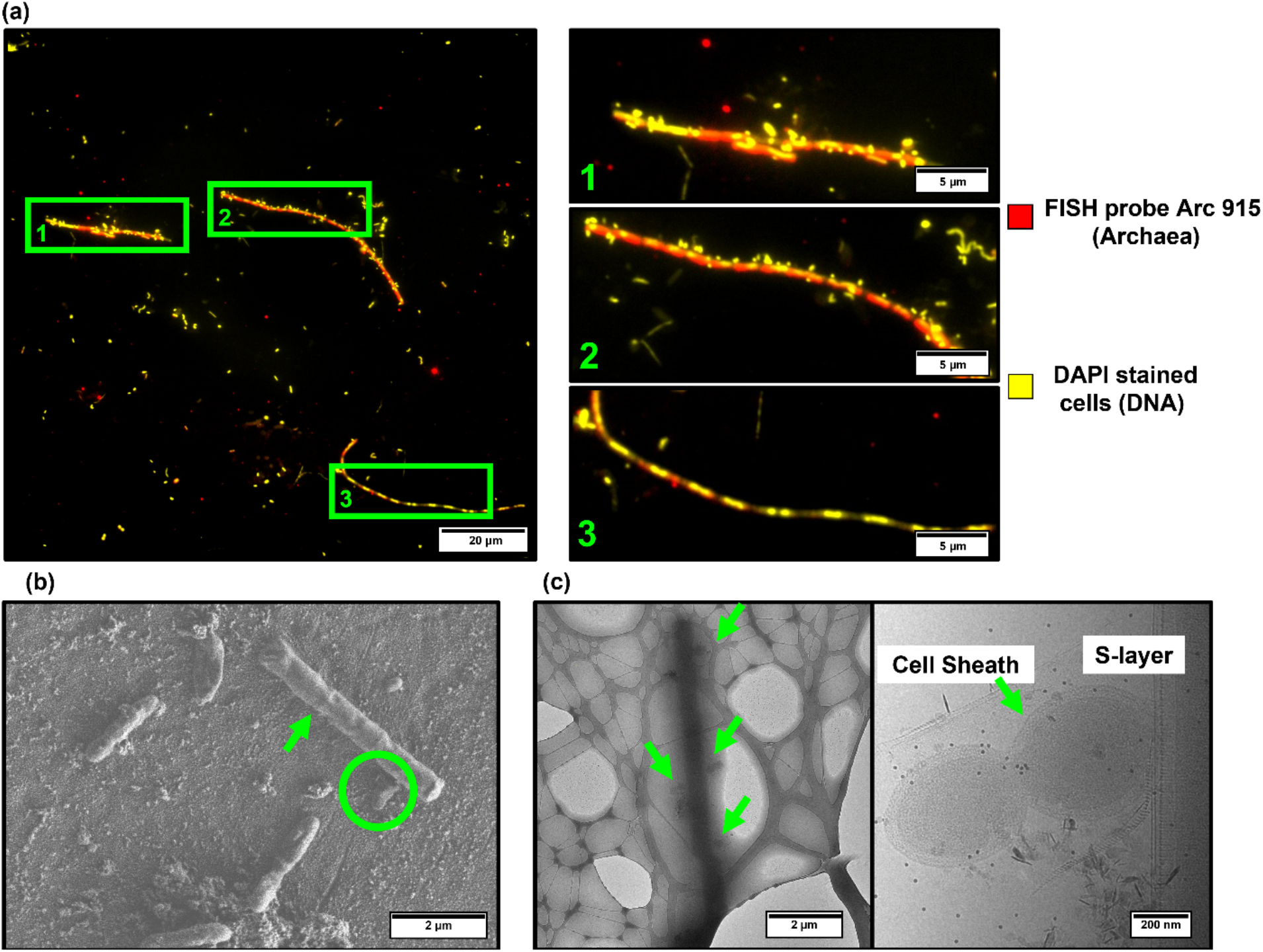
Various microscopic images of *Ca*. Nealsonbacteria in enrichment cultures. (a) Epifluorescence microscopy showing archaea (red) and DAPI (yellow). Images are false colored and overlaid using ImageJ. Images on the right are zoomed in views of green rectangular area in the left image highlighting archaeal filaments with and without attached cells. (b) SEM images of methanogenic benzene-degrading culture prepared by ionic liquid exchange. A *Methanothrix* filament is marked with green arrow. *Ca*. Nealsonbacteria-like cells are marked with a green circle. (c) Cryto-TEM images of *Ca*. Nealsonbacteria enrichments. *Ca*. Nealsonbacteria-like cells are marked by green arrows and the right-side shows a zoomed in view of two cells that appear to be budding. Additional microscopic images are shown in Figures S5 and S6.

We had previously unsuccessfully attempted to enrich *Ca*. Nealsonbacteria by filtration through a 0.22-μm filter. Analysis of the filtrate by qPCR showed no enrichment: both bacterial and the *Ca*. Nealsonbacteria concentrations decreased 100-fold (data not shown). Other studies have reported success enriching ultra-small microbes this way (1, 26, 28). Perhaps the strong attachment of *Ca*. Nealsonbacteria to its host caused this approach to fail. Interestingly, we also noticed that the archaeal FISH probe seemed to bind more strongly to methanogens that had many *Ca*. Nealsonbacteria cells attached, compared to naked methanogens (Figure 4a, bottom right; Figure S5). Naked methanogens stained brightly with DAPI (DNA) but only faintly with the archaeal FISH probe (rRNA). Given that FISH signal intensity is a measure of ribosome count and thus metabolic activity (29), perhaps methanogens with *Ca*. Nealsonbacteria attached are more active. This observation is in contrast to the recent study by Kuroda et al., (27) where a major fraction of the Ca. Nealsonbacteria were attached to methanogens with weaker fluorescence. *Ca*. Nealsonbacteria’s role as either a parasitic or commensal organism is thus still unclear.

### SEM and Cryo-EM confirm attachment to *Methanothrix*

SEM examination of the parent methanogenic benzene-degrading culture prepared with ionic liquid exchange to preserve cell structure also showed a similar association of very small *Ca*. Nealsonbacteria-like cells attached to a large filamentous cell structure consisting of a chain of rod-shaped, blunt-ended cells (Figure 4b). This filament was enclosed in a tubular sheath and the flat ends distinctively match the characteristics of *Methanothrix* (30). The *Ca*. Nealsonbacteria-like cells are spherical with a diameter of 0.2 μm to 0.8 μm and they were always found in budding pairs, forming a peanut shape. Cryo-EM was also used to examine the *Ca*. Nealsonbacteria enrichments. The Cryo-EM images (Figure 4c) show the same association of peanut-shaped cells with *Methanothrix*. The large black rod-shaped blurry cells at the center of the image are *Methanothrix* forming a sheathed chain. Multiple peanut-shaped ultra-small cells were observed attached to the chain of *Methanothrix* (Figure 4c). The *Ca*. Nealsonbacteria-like cells were always found in pairs and the one attached to the methanogen was usually bigger. A zoomed-in image (Figure 4c; right side) reveals that *Ca*. Nealsonbacteria cells have a very interesting membrane structure with a thick sheath and an apparent S-layer. Additional supporting FISH, SEM, Cryo-EM images are provided in Figures S5 and S6. The close association of *Ca*. Nealsonbacteria and *Methanothrix* may facilitate efficient nutrient exchange between these organisms. Given that *Methanothrix* are obligate acetoclastic methanogens, *Ca*. Nealsonbacteria may provide acetate to *Methanothrix*. The physical association also enables potential direct interspecies electron transfer or DIET (31). Although, e-pili genes were not found in the *Ca*. Nealsonbacteria genome (see below), DIET or direct acetate exchange could happen by other mechanisms. Clearly the possible syntrophy between the *Ca*. Nealsonbacteria and *Methanothrix* requires further investigation.

### A co-occurrence network finds *Ca*. Nealsonbacteria variants positively correlated with *Methanothrix*

A co-occurrence network inference (CoNet) analysis was conducted using all 16S rRNA amplicon sequencing data from the two donor trials (Figure S7a). *Ca*. Nealsonbacteria ASVa was found to correlate positively with its putative host *Methanothrix* ASV89, as well as to *Ca*. Nealsonbacteria ASVb and to two less abundant microbes (*Sporomusa* and *Ignavibacteriales*) and to correlate negatively with *Spirochaetaceae* ASV977, which was only enriched in the bottles amended with pyruvate. *Ca*. Nealsonbacteria ASVb (enriched only when pyruvate or acetate were added) was positively correlated with *Methanothrix* ASV89 as well as to *Methanothrix* ASV91. ASV91 shares 99.4% pairwise identity with *Methanothrix* ASV89 and could be a specific host for *Ca*. Nealsonbacteria ASVb. A heatmap of absolute abundance was generated (Figure S7b) that includes all the ASVs from the co-occurrence network. The rows of heatmap were clustered based on similarity. Interestingly, *Ca*. Nealsonbacteria ASVa clustered with *Methanothrix* ASV89 while *Ca*. Nealsonbacteria ASVb clustered with *Methanothrix* ASV91 (Figure S7b, red rectangles). This clustering suggests that *Ca*. Nealsonbacteria ASVa and ASVb may have different *Methanothrix* hosts.

### Impact of *Ca*. Nealsonbacteria on benzene degradation

To further study the role of *Ca*. Nealsonbacteria in benzene degradation, we mixed an enrichment (containing about 10^7^ *Ca*. Nealsonbacteria cells/mL) with a “slow” benzene-degrading culture (containing about 10^7^ ORM2 cells/mL, degrading at ~0.1 mg/L/day). As a control, we also amended the slow culture with supernatant recovered from a healthy culture (degrading benzene at 10 mg/L/day). The raw data from this experiment are reported in Table S8. We observed a significant increase in benzene degradation rate in the two replicates that were a 50:50 mix of the two cultures (Figure 5a), even though mixing diluted the benzene degrader (ORM2) by 50%. Therefore, adding a *Ca*. Nealsonbacteria enrichment enhanced benzene degradation. Slow benzene culture amended with supernatant from a healthy culture as well as the slow culture-only controls all showed overall slower benzene degradation (Figure 5a). Benzene biodegradation rates were highest in cultures with the highest measured ORM2 concentrations at Day 217, except for one culture replicate (slow culture #1) with abundant ORM2 but no benzene degradation activity (Figure 5b). The two vials amended with the *Ca*. Nealsonbacteria enrichment showed the highest cell concentrations for both ORM2 and *Ca*. Nealsonbacteria. Perhaps *Ca*. Nealsonbacteria supplied ORM2 or *Methanothrix* with specific limiting metabolites, nutrients or cofactors recycled from lysed biomass or simply removed inhibitory compounds. Further studies using isotope-labeled cell lysate may help to clarify these complex interactions.

**Figure 5.**
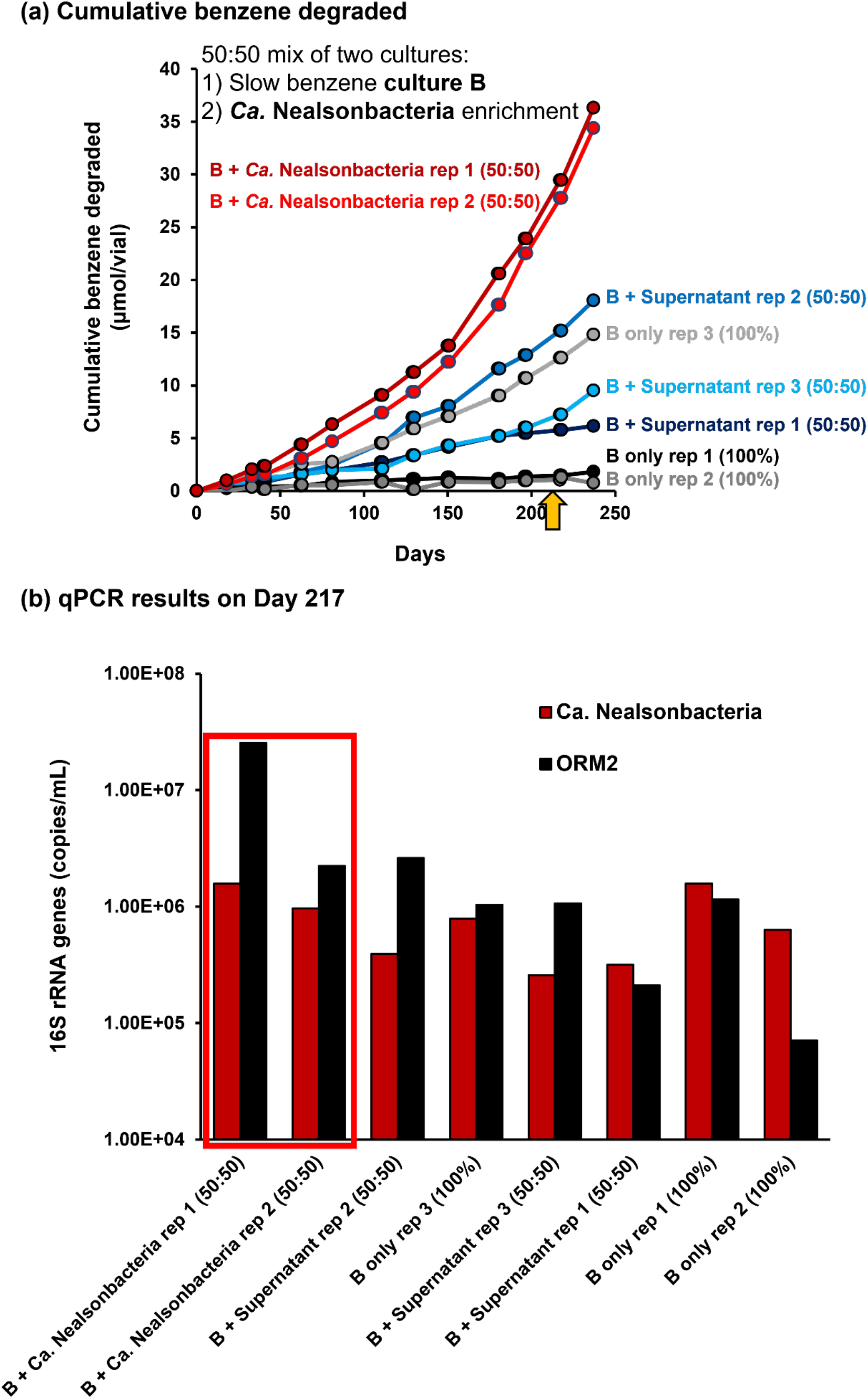
Impact of adding *Ca*. Nealsonbacteria enrichment to a slow benzene-degrading culture. a) Cumulative benzene consumption and b) qPCR results tracking *Ca*. Nealsonbacteria and ORM2 on Day 217. The qPCR results are sorted by descending order of benzene degradation rate. The biological replicates are plotted separately to show the differences.

### The genome of *Ca*. Nealsonbacteria DGGOD1a is consistent with a symbiotic epibiotic lifestyle

We assembled and closed the genome of *Ca*. Nealsonbacteria DGGOD1a (Genbank: CP092821 and IMG ID: 2791354853) corresponding to ASVa using both Illumina and PacBio sequence data. The details of assembly are summarized in Supplemental Text S4 and reported in a genome announcement (55). The quality of the genome assembly was validated with read-mapping and a GC skew plot (Figure S8) clearly showing origin and terminus. The complete genome is 1.16 Mbp in length with 1173 predicted coding sequences (CDS) including 18 pairs of perfect repeats which are over 400 bp in length. The majority of these repeats are predicted transposases, and in fact, this genome has a total of 39 predicted transposases which comprise 3.4% of protein coding genes.

Transposases are involved in genomic rearrangements, gene duplication and for promoting horizontal gene transfer (32). A total of nine regions were predicted as genomic islands using Islandviewer4 (31) of which seven contain transposases (Table S9). High numbers of transposases are found in organisms under stress or in extreme environments (32) which may explain the unusual high decay rate sometimes seen in our methanogenic benzene degrading culture (19). Other studies have also shown a high transposase content in bacteria with symbiotic or pathogenic lifestyles (33–35). The transposases in the genome of *Ca*. Nealsonbacteria may participate in DNA exchange with the host *Methanothrix*.

Like other CPR bacteria, the genome reveals that most metabolic pathways and biosynthetic capacities annotated by automatic pipelines are predicted to be incomplete, notably of nucleotides, lipids, vitamins and most amino acids (Figure 6). As inferred from genomes of other members of the CPR group (1, 5, 7), DGGOD1a can use the pentose phosphate pathway to bypass glycolysis. Also, there is no lipid biosynthesis pathway and the pathways for the biosynthesis of most amino acids and vitamins are incomplete. The only vitamin DGGOD1a is predicted to make is riboflavin. The large ribosomal RNA subunit L1 is missing; a common feature of members of the OD1-L1 group (1, 5). Interestingly, eight toxin and five antitoxin gene families were found in the genome. Some were found located within two predicted genomic islands and may contribute to their maintenance (36). The toxin-antitoxin systems are stress-response elements that could also help cells adapt to unfavorable growth conditions (36). These systems are common in free-living microbes but tend to be lost from host-associated prokaryotes due to the relatively constant environment of host-associated organisms (37). The high toxin-antitoxin genes copies in *Ca*. Nealsonbacteria indicates that it is possibly still undergoing reductive evolution (36, 38). Twelve type IV pilus assembly proteins, two cell membrane proteins annotated as holins, and type II secretion system genes were found in the genome, which are consistent with an epibiont lifestyle. Epibionts are known to use type IV pili to attach to the host membrane (39). The holin protein family form pores in cytoplasmic membranes and may be used to permeabilize the host cell membrane and perhaps kill the host (40). Type II secretion systems may be involved in nutrient uptake or nutrient exchange between the *Ca*. Nealsonbacteria and its host and may also participate in DIET (41). Surprisingly, a type V CRISPR-Cas system that contains 49 spacers was found in the genome with Cas12a as the effector and the complex of cas4, cas1 and cas2 as the spacer acquisition machinery. It is rare to see a CRISPR-Cas system in such a reduced genome, and only 2.4% of organisms from the Parcubacteria and Microgenomates superphyla have a CRISPR-Cas system (42). The intact CRISPR-Cas system suggests exposure to phage and extracellular DNA. Considering that *Ca*. Nealsonbacteria lack the ability to synthesize nucleotides, perhaps the CRISPR-Cas system is involved in the degradation of host DNA for recycling nucleotides. No clues were gleaned from a search of CRISPR spacer targets against the OR consortium metagenome and all the sequences in the NCBI database, as these searches only returned hits to the CRISPR matrix itself.

**Figure 6.**
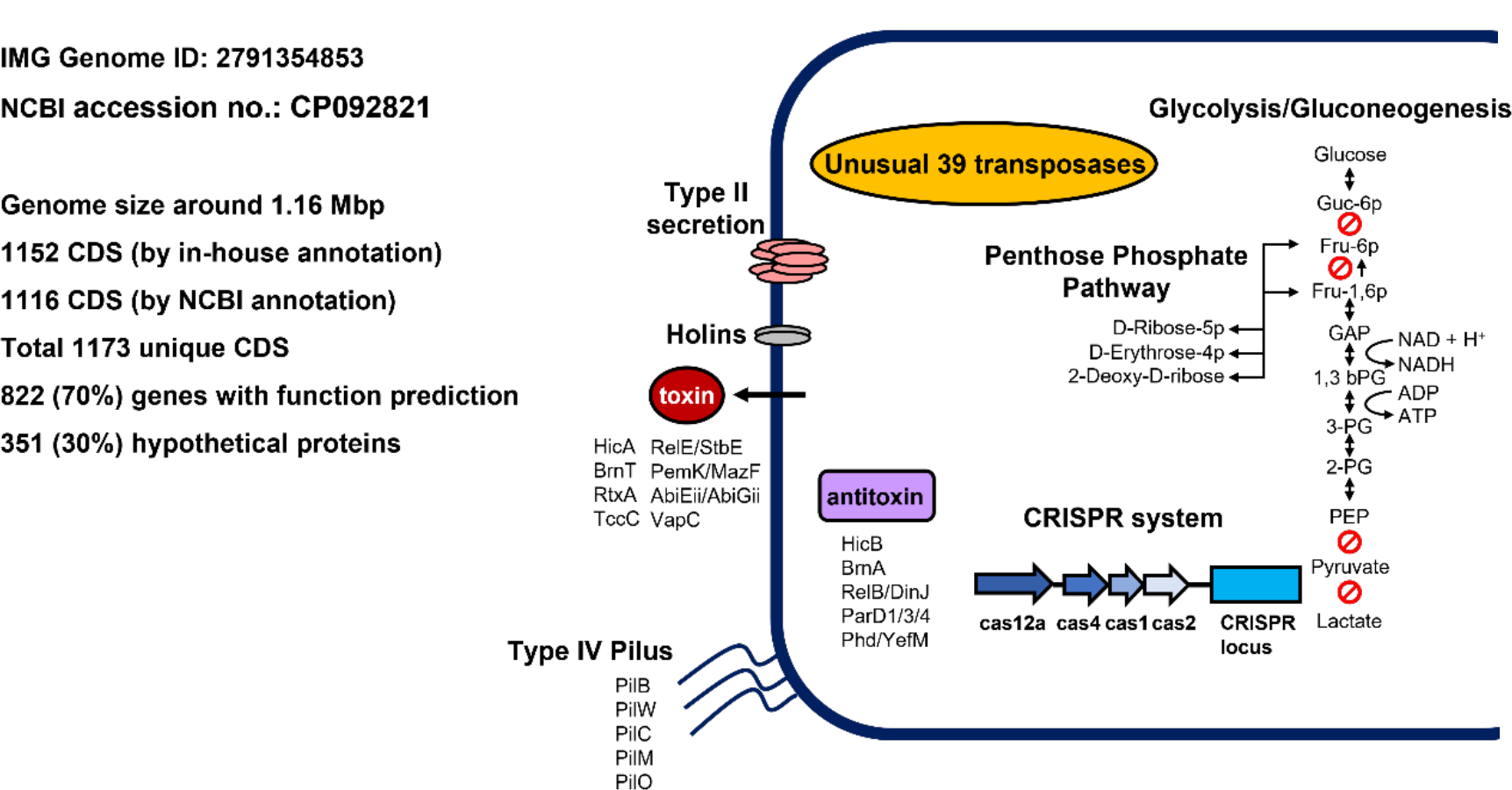
A summary of key findings from the genome of *Ca*. Nealsonbacteria DGGOD1a. Detailed annotations for each ORF are in Table S10.

We were not able to assign function to a large proportion (30%) of the predicted proteins in the genome, despite running multiple prediction pipelines and manual curation (Table S10). One interesting feature is a putative NADP-reducing hydrogenase. This protein has been found in the bacterium *Desulfovibrio fructosovorans* as an iron-sulfur protein that exclusively functions as a hydrogen dehydrogenase (43) and may mean that Ca. Nealsonbacteria uses hydrogen produced by other fermentation reactions. The genome also encoded two putative polysaccharide deacetylases belonging to carbohydrate esterase family 4 (CE4). The members of this family can hydrolyze the acetyl group from *N*-acetylglucosamine (44–47) or *O*-acetylxylose residues (48, 49) which exist in extracellular polymeric substances (EPS). The released acetyl group could be used by the obligate acetoclastic methanogenic archaeon *Methanothrix*. Three genes encoded putative peptidoglycan-binding proteins were also found in the genome. The proteins belong to peptidoglycan-binding domain could involve in cell wall synthesis (50) and binding (51) which may responsible for the complex membrane structure we found by Cryo-EM.

### Implications

*Ca*. Nealsonbacteria DGG1a likely contributes to recycling biomass including extracellular polymeric substances in a methanogenic benzene degrading enrichment culture, and in so doing enhances the growth of benzene fermenters. A role in microbial biomass recycling is supported by substantial enrichment of this phylotype on lysed biomass. The finding of a close association of dividing *Ca*. Nealsonbacteria cells with *Methanothrix* is now the second report of a CPR bacterium that is an episymbiont of an archaeon (27), in this study shown in a stable laboratory enrichment culture where further experimentation is possible. *Ca*. Nealsonbacteria were observed to be beneficial for the growth of benzene-metabolizing *Desulfobacterota* ORM2 from yet another uncultivated lineage. *Ca*. Nealsonbacteria may recycle key cofactors required to initiate benzene ring activation. These results guide future experiments into specific relationships between *Ca*. Nealsonbacteria and benzene degradation in these enrichment cultures.

The visualization of this unusual cross-kingdom episymbiosis of *Ca*. Nealsonbacteria on the surface of *Methanothrix* implies metabolite exchange or direct electron transfer between the two cell types. The type IV pili encoded in the genomes of many *Ca*. Nealsonbacteria (5, 7, 52) are possibly used for attachment to the host and as well as a conduit for electron transfer. Perhaps these tiny microbes revealed in many metagenomic surveys of natural and engineered anaerobic communities have analogous roles? *Ca*. Nealsonbacteria may be particularly relevant where energy input is limited and recycling is vital, such as in deep subsurface environments.

## Experimental Procedures

### Chemicals and microbial cultures

All chemicals were obtained from Sigma-Aldrich (Oakville, ON, Canada) at the highest purity available unless specified otherwise. The OR consortium is maintained in 100 mL to 2 L batch culture bottles at the University of Toronto and in larger vessels (>100 L) at SiREM laboratories (Guelph, Ontario, Canada); the scaled-up culture is referred to as DGG-B (17). Two OR consortium subculture bottles (or lineages) were used in the experiments described herein: a bottle called OR-1bBig was the inoculum for *Ca*. Nealsonbacteria enrichment Donor Trials #1 & #2 and a bottle called OR-p5 was the inoculum for the augmentation experiment. All cultures were grown in an iron sulfide (FeS)-reduced, bicarbonate-based mineral medium (MM medium) amended regularly (approx. 1 per month) with benzene targeting aqueous concentration of 10 mg/L per bottle (17, 18).

### Preparation of electron donor solutions for *Ca*. Nealsonbacteria enrichment trials

Two crude cell lysate solutions from the OR consortium OR-1bBig and a culture of *Escherichia coli* were prepared using tangential flow filtration and French press as described in Text S1. Next, we prepared two solutions of specific biomass components, nucleic acids (salmon sperm DNA; Sigma-Aldrich, Deoxyribonucleic acid sodium salt) and a membrane lipid (L-α-phosphatidylethanolamine; Sigma-Aldrich, Type II, ≥97%, lyophilized powder from egg yolk). Acetate, pyruvate and hydrogen were also tested as donors as they are intermediates of benzene degradation. Stock solution concentrations are listed in Table S1a.

### *Ca*. Nealsonbacteria Enrichment – Donor Trial #1

To test if *Ca*. Nealsonbacteria DGGOD1a could grow on crude lysed biomass or specific sub-cellular components, 25 mL culture tubes with 9 mL MM medium were inoculated with 1 mL OR-1bBig culture and amended with 7 different electron donor solutions (Table 14a) at approximately the same electron equivalents, targeting ~30 mg COD/L. Each condition was tested in quadruplicate and incubated in a Coy anaerobic glove box for 7 months (supplied with 10% H_2_, 10% CO_2_, and 80% N_2_). DNA samples were collected at 2, 4, and 7 months after inoculation by sacrificing an entire replicate from each treatment.

### *Ca*. Nealsonbacteria enrichment – Donor Trial #2

After observing successful growth and enrichment of *Ca*. Nealsonbacteria with crude cell lysate from the benzene culture in Donor Trial #1, a second experiment was set up to test the impact of various biomass fractions of cell lysate. Culture (500 mL) was concentrated by tangential flow and cells were lysed using the same French press method described earlier (Text S1). The cell lysate was then divided into four fractions, as seen in Figure S3. One fraction was used “as is” (BL for <u>B</u>enzene culture <u>L</u>ysate). The remaining lysate was centrifuged (at 13,000 × g for 20 mins) to create a pellet fraction and a supernatant fraction. The supernatant fraction was further separated by tangential flow filtration with 50kDa cut-off membrane (Pellicon® XL cassette with Durapore® membrane) to create a permeate fraction (< 50 kDa) and a retentate fraction (>50 kDa). Each of these fractions was tested separately (Table S1b). Two additional treatments were included in Donor Trial #2. One consisted of pyruvate plus the antibiotics kanamycin and vancomycin (1 mM) to inhibit bacteria without inhibiting methanogens. A second additional treatment was amended with *E. coli* extract instead of benzene culture extract. We ensured that the *E. coli* pellet was well washed with MM medium to remove any residual *E. coli* rich growth medium that could confound results. Controls without substrate (starved control; SC) and crude benzene culture lysate without inoculum (no inoculum; NI) were included. All treatments were conducted in biological duplicates except the starved control which was a singleton.

### *Ca*. Nealsonbacteria augmentation trial

This experiment made use of an inoculum from culture bottle “OR-p5” with a very slow benzene degradation rate over years (~0.1 mg/L/day). Eight 25-mL culture vials were each filled with a 10 mL aliquot of this slow culture (see Text S2 for details). Subsequently, 5 mL of OR-p5 culture was removed from vials #1-2 and replaced with 5 mL of *Ca*. Nealsonbacteria enrichment culture BL (replicate #1) from Donor Trial #2. In vials #3-5, 5 mL of culture was removed and replaced with 5 mL of culture supernatant from a very active benzene culture (degradation rate of ~10 mg/L/day), prepared by centrifugation at 13,000 × g for 20 min. The remaining vials (#6-8) of slow culture were left as is. All vials were amended with neat benzene targeting 15 mg/L aqueous concentration and reamended whenever concentrations decreased below 5 mg/L. Benzene and methane concentrations were monitored by GC/FID (see below). DNA samples were collected on day 217 for qPCR analysis.

### DNA extraction, quantitative PCR, and Illumina amplicon sequencing and statistical analysis

DNA from Donor Trial #1 was extracted from 10 mL culture samples; otherwise, DNA was extracted from 1 mL culture samples. Cells were harvested by centrifugation at 13,000 × g for 20 min at 4°C and pellets resuspended in residual (~0.05 mL) supernatant. DNA was extracted using the DNeasy PowerSoil Kit (QIAGEN) following the manufacturer’s protocol. Real-time quantitative polymerase chain reaction (qPCR) assays were performed to track the gene copy numbers of Bacteria and Archaea using universal 16S rRNA gene primers; abundances of *Ca*. Nealsonbacteria and *Thermodesulfobacteria* ORM2 were also tracked using specific 16S rRNA gene-targeting primers (Table S11). qPCR reactions were performed using a CFX96 real-time thermal cycler (Bio-Rad Laboratories) using the following thermocycling conditions: an initial denaturation step at 98 °C for 2 min, 40 cycles of 98 °C for 5 s and Tm (see Table S11) for 10 s, followed by melt curve analysis (65-95 °C with an increase of 0.5 °C every 5 s). qPCR results were processed with CFX Manager software (Bio-Rad Laboratories). Standard curves for all the qPCR studies are reported in Table S12. 16S rRNA gene amplicon sequencing with primers targeting the V6-V8 region was conducted at the Genome Quebec Innovation Centre (McGill University) as previously described (17). Raw reads were processed into amplicon sequence variants (ASVs) and analyzed by nonmetric dimensional scaling (NMDS) and Co-occurrence Network Inference (CoNet) analysis (53) as described in Text S3.

### Metagenome analysis

The OR consortium/DGG culture was sequenced using a combination of Illumina paired-end (~150 bp reads, ~300 bp insert) and PacBio long-read library (~7000 bp reads). Metagenomes were submitted to IMG under sequencing project Gp0324998. Detailed description of assembly, binning and functional and taxonomic annotation can be found in Text S4.

### Epifluorescence microscopy and fluorescence in situ hybridization (FISH)

The FISH probe ARCH915 was used for visualizing all Archaea (54). Eight FISH probes (Table S6) targeting the 16S rRNA gene of *Ca*. Nealsonbacteria were designed and tested in this study but none yielded a signal distinguishable from background noise. The protocol for epifluorescence microscopy is provided in Text S5 and details into the design and testing of FISH probes in Text S6.

### Scanning Electron Microscopy with Ionic Liquid and Cryo-EM

To maintain the wet native state of microbial cells and to eliminate artifacts that may be introduced by sample drying and coating, culture samples were prepared with an ionic liquid and examined by SEM, as detailed in Text S7. Cryo-EM was also used to explore intracellular structure and spatial associations. Sample preparation and Cryo-EM conditions are described in Text S8.

### Analytical procedures

Benzene and methane were measured in culture bottles and vials by injecting 0.3 mL headspace samples into a Hewlett-Packard 5890 Series II GC fitted with a GSQ column (30 m x 0.53 mm I.D. PLOT column; J&W Scientific, Folsom, CA) as previously described (18). External standards were used for calibration.

### Accession numbers

The closed genome sequence of *Candidatus* Nealsonbacteria DGGOD1a is available at the Joint Genome Institute (IMG) under Taxon ID 2791354853 and at NCBI under accession no. CP092821. Amplicon sequences (16S rRNA) were uploaded to the NCBI SRA under BioProject accession number PRJNA830497. The Genome is also available in ggkbase with the following link: https://ggkbase.berkeley.edu/organisms/410827/contigs/251537015.

## Supporting information

Supplemental large Tables

## Supporting information

The PDF file contains: 1) supplementary Figures S1 to S6, including a phylogenomic tree of *Ca*. Nealsonbacteria, fractionation steps used to preparer biomass, NDMS plots, *Ca*. Nealsonbacteria growth curves and additional microscopic images; 2) supporting texts (S1 to S8) providing additional method details; and 3) supplementary Tables S9, S10, S11 listing primers, probes and qPCR standard curves. The excel file contains larger data Tables S1 and S5-S12.

## Acknowledgements

This study was funded by Genomic Application Partnership Program (GAPP) grants awarded to E.A.E. (Project IDs OGI-102 and OGI-173), which were supported by Genome Canada, Ontario Genomics, the Government of Ontario, Mitacs Canada, SiREM, Alberta Innovates, Federated Co-operatives Limited, and Imperial Oil. The authors would like to thank SiREM (Guelph, Ontario N1G 3Z2, Canada) for supplying culture for testing. We would also like to acknowledge the laboratories of Dr. Neil Thomson (University of Waterloo), Dr. Ania Ulrich (University of Alberta), SiREM, and Innotech Alberta (Edmonton, Alberta T6N 1E4, Canada) for providing feedback and intellectual contributions to this study during weekly GAPP consortium meetings.

## AUTHOR CONTRIBUTIONS

All authors made research and substantial intellectual contributions to the completion of this study. X.C. and E.A.E. conceived and designed the study, and X.C carried out all experiments, data analysis and interpretation, and drafted and revised the manuscript. O.M. assembled the OD1 genome with help of C.N. and C.T.B. L.M performed the Cryo-TEM data acquisition and analysis. C.R.A.T. helped with data analysis and revised the manuscript. S.G. helped to maintain the culture, performed qPCR data acquisition and analysis and revised the manuscript. F.L. enriched the culture, extracted DNA and performed the initial metagenome analysis. J.H. assisted with ionic liquid-SEM data acquisition and analysis. C.H. performed the OD1 taxonomy classification, phylogenetic tree and OD1 genome annotation. J.H.D.C. and J.F.B. provided critical research guidance, and manuscript revisions.

