## Supplemental information for "*Candidatus* Nealsonbacteria (OD1) are biomass recycling ectosymbionts of methanogenic archaea in a stable benzene-degrading enrichment culture"

Number of Pages: 22

Number of SI Figures: 8

Number of Supporting Texts: 8

Number of SI Tables: 12

**TABLE OF CONTENTS**

**Supplementary Figures**

**Figure S1:** Phylogenomic tree of the *Ca.* Nealsonbacteria found in the OR/DGG-B culture and its closest relatives
**Figure S2:** Non-metric dimensional scaling analysis of *Ca.* Nealsonbacteria Donor Trials #1 & #2 based on relative abundances of each ASV
**Figure S3:** Detail fractionation steps for *Ca.* Nealsonbacteria enrichment biomass fractionation trial
**Figure S4:** *Ca.* Nealsonbacteria growth curve for biomass fractionation donor Trial #2
**Figure S5:** Additional FISH images for *Ca.* Nealsonbacteria enrichment culture
**Figure S6:** Additional SEM images for methanogenic benzene-degrading culture (top row) and Cryo-EM images for *Ca.* Nealsonbacteria enrichment culture (bottom row)
**Figure S7:** Network interactions between Ca. Nealsonbacteria ASVa and ASVb and their correlated microbes (a) and corresponding heatmap (b) based on absolute abundance.
**Figure S8:** GC skew and cumulative GC skew of *Ca.* Nealsonbacteria DGGOD1a genome.

**Supporting Texts**

**Text S1:** Culture lysate preparation for Donor Trials
**Text S2:** Sample pretreatment for *Ca.* Nealsonbacteria augmentation trial
**Text S3:** 16S rRNA gene amplicon raw read processing, NMDS and CoNet analysis.
**Text S4:** Metagenomic assembly, binning and functional and taxonomic annotation
**Text S5:** Epifluorescence microscopy
**Text S6:** Fluorescence in situ hybridization (FISH) method details
**Text S7:** Ionic liquid sample preparation and SEM setup
**Text S8:** Cryo-EM sample preparation and operational condition

**Supplementary Tables (larger tables are in accompanying Excel file)**

**Table S1:** Treatment tables for Donor Trials #1 and #2 (**Excel**)
**Table S2:** qPCR raw data for Donor Trial #1 and #2 (**Excel**)
**Table S3:** List of all gDNA samples amplified by illumina sequencing and their taxonomic assignment (**Excel**)
**Table S4:** Methane production and electron balance in Donor Trial #1 and methane production in Donor Trial #2 (**Excel**)
**Table S5:** Raw data used to plot the Heatmap of Donor Trial #2 (**Excel**)
**Table S6:** Specific FISH probes used in this study
**Table S7:** Cell count raw data for epibiont small cells in *Ca.* Nealsonbacteria Enrichment (**Excel**)
**Table S8:** Raw data for *Ca.* Nealsonbacteria Augmentation Trial (**Excel**)
**Table S9:** Possible genome Islands predicted by Islandviewer 4 (**Excel**)
**Table S10:** Genome annotation determined by hidden Markov models search against KEGG, Uniref, TIGRfam, Pfam, Custom and NCBI annotation pipeline (**Excel**)
**Table S11:** Specific 16S rRNA primers used for qPCR in this study
**Table S12:** Summary of all qPCR standard curve equations and efficiencies

**References for SI**

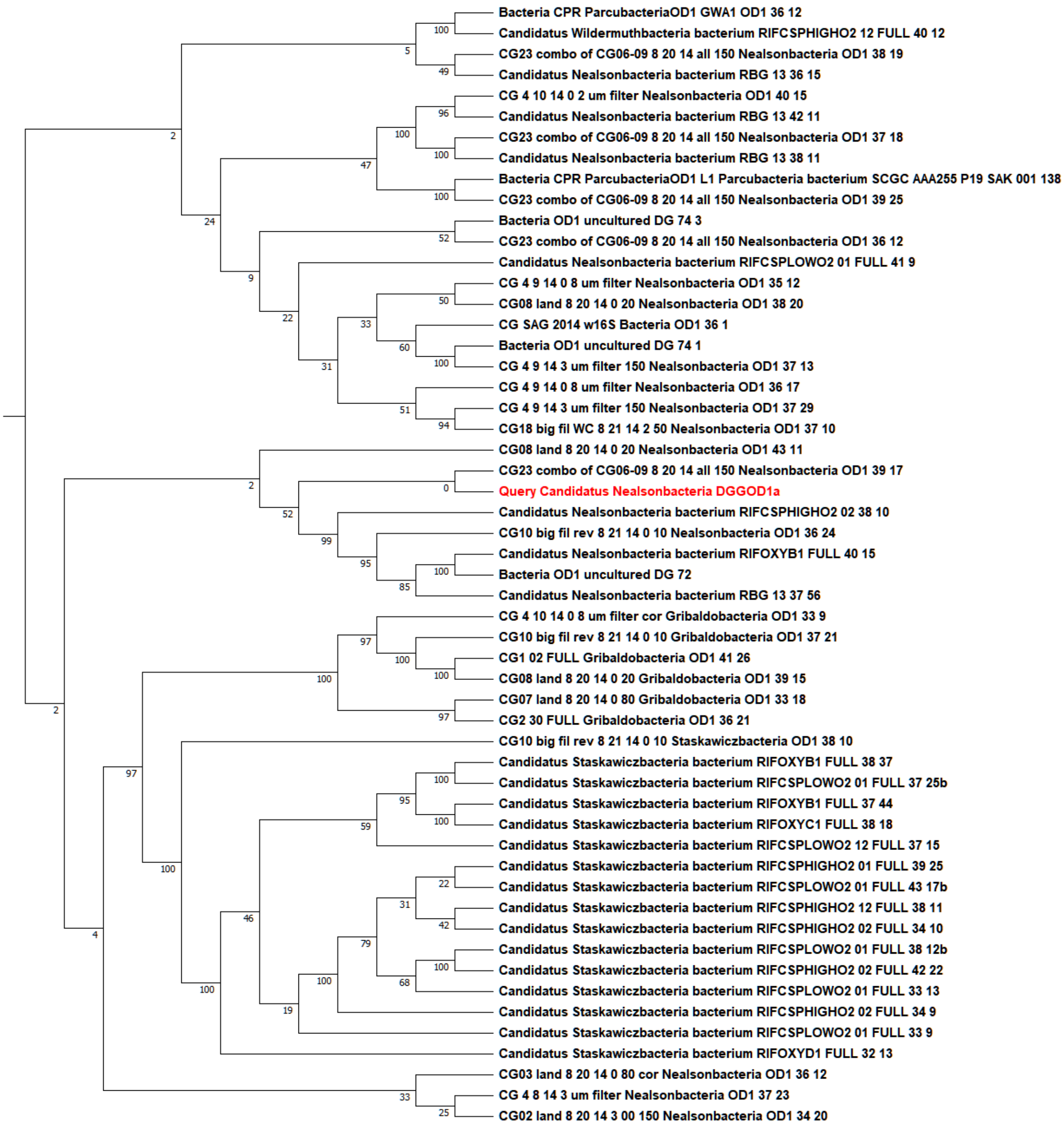


**Figure S1: Phylogenomic tree of the *Ca.* Nealsonbacteria found in the OR/DGG-B culture and its closest relatives**. The studied microbe, shown in red, is named "*Candidatus* Nealsonbacteria DGGOD1a". The figure presents a subtree of a maximum likelihood concatenated ribosomal protein tree with references from across bacteria and archaea, including a large number of previously published high-quality CPR and DPANN genomes. Tree construction is described in Text S4.

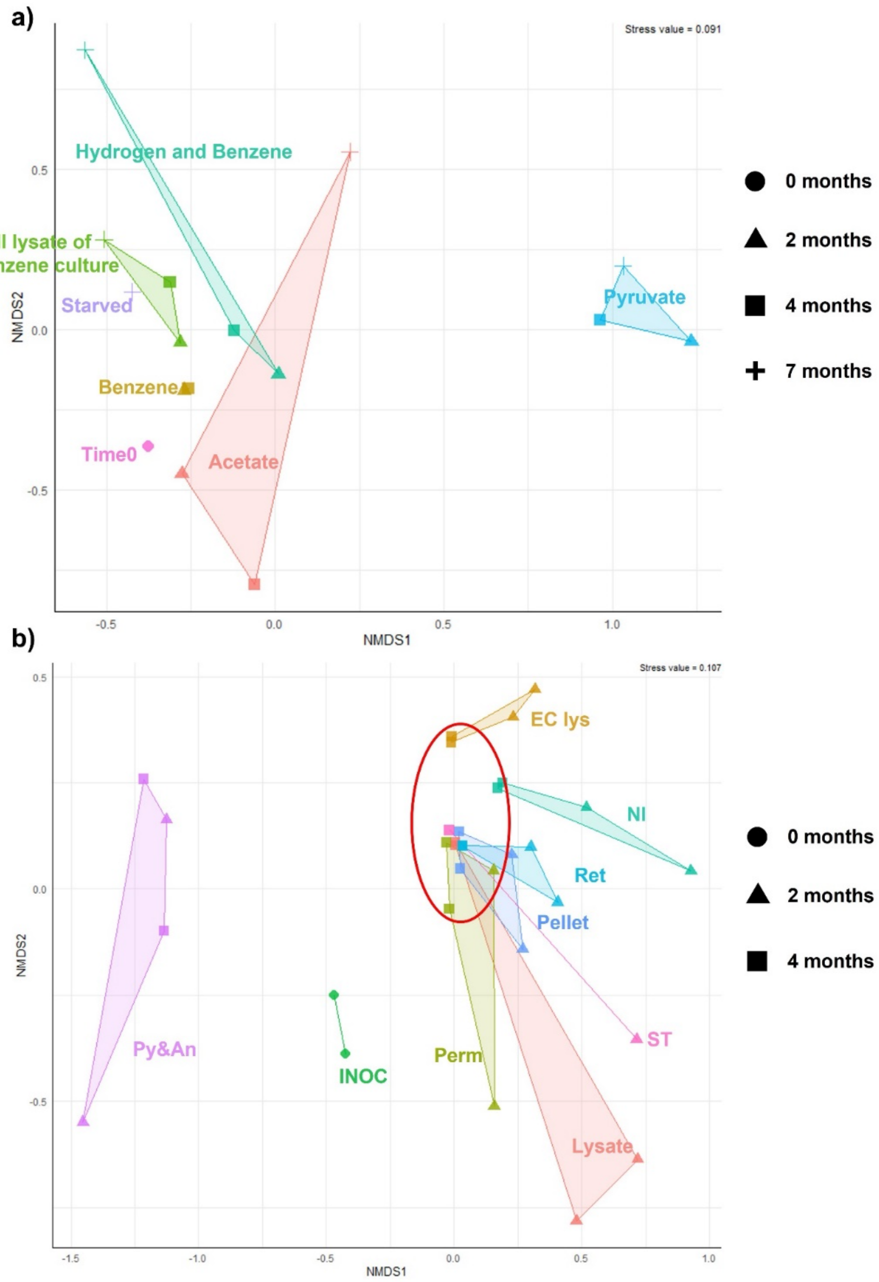


**Figure S2: Non-metric dimensional scaling analysis of *Ca.* Nealsonbacteria Donor Trial #1 (Panel a) and Donor Trial #2 (Panel b) based on relative abundances of each ASV**. Sample points are grouped by different treatment by colour. The shapes (circles, triangles, squares) correspond to different incubation times, and the red circle highlights convergence of data after 4 months (squares). The line connecting all the sample points helps to locate points in each group. The group names coded as follows: INOC: Bottle with only 10% inoculum at time 0; NI: no inoculum (lysate only); Py&An: pyruvate & antibiotics; Lysate: French pressed benzene culture; Perm: lysate supernatant permeate through 50 kDa membrane; Ret: lysate supernatant retentate; Pellet: lysate pellet after centrifugation at 13,000 × g for 20 min; EC lys: French press lysate of *E. coli* culture; ST: Starved culture.

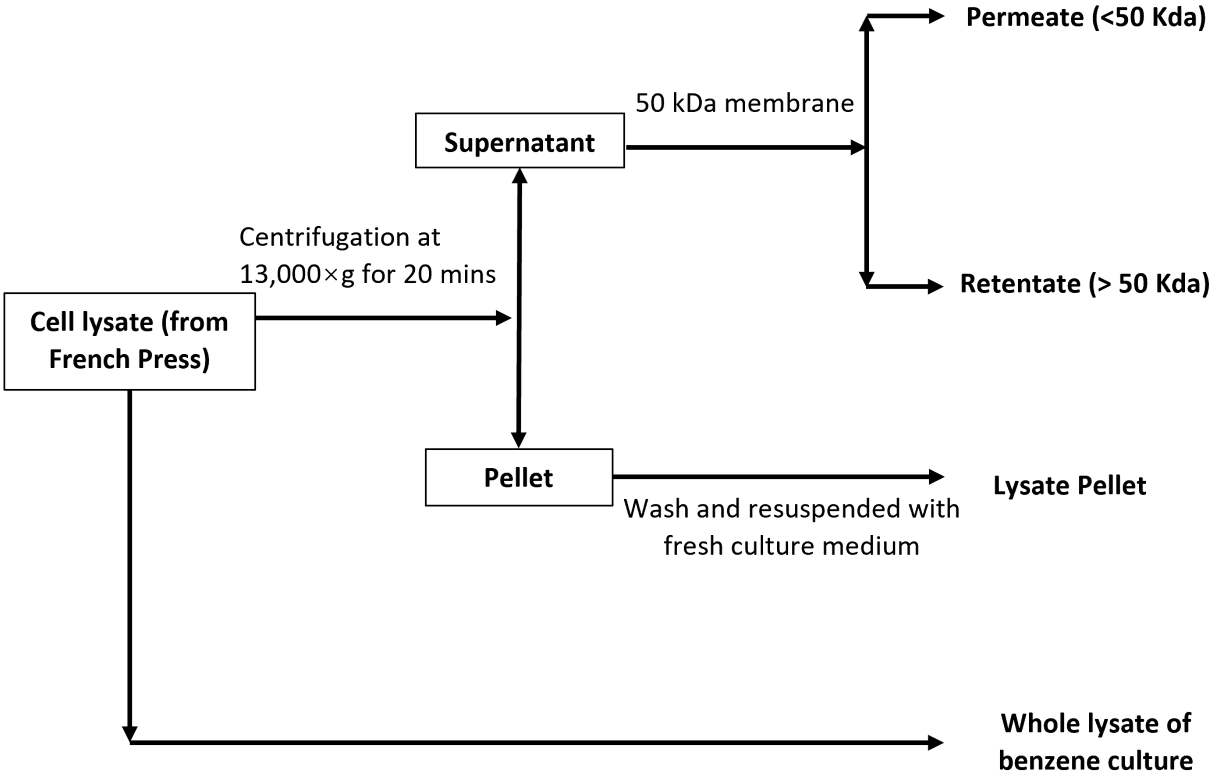


**Figure S3: Detail fractionation steps for *Ca.* Nealsonbacteria enrichment biomass fractionation trial #2.** Each treatment was carried out in biological duplicates. Details for bottle setup are provided in Table S1b.

Codes used for each fraction:
**Perm:** Lysate supernatant permeate (< 50 kDa)
**Ret:** Lysate supernatant retentate (> 50 kDa)
**Pellet:** Lysate pellet
**Lysate:** French Press Lysate of benzene culture

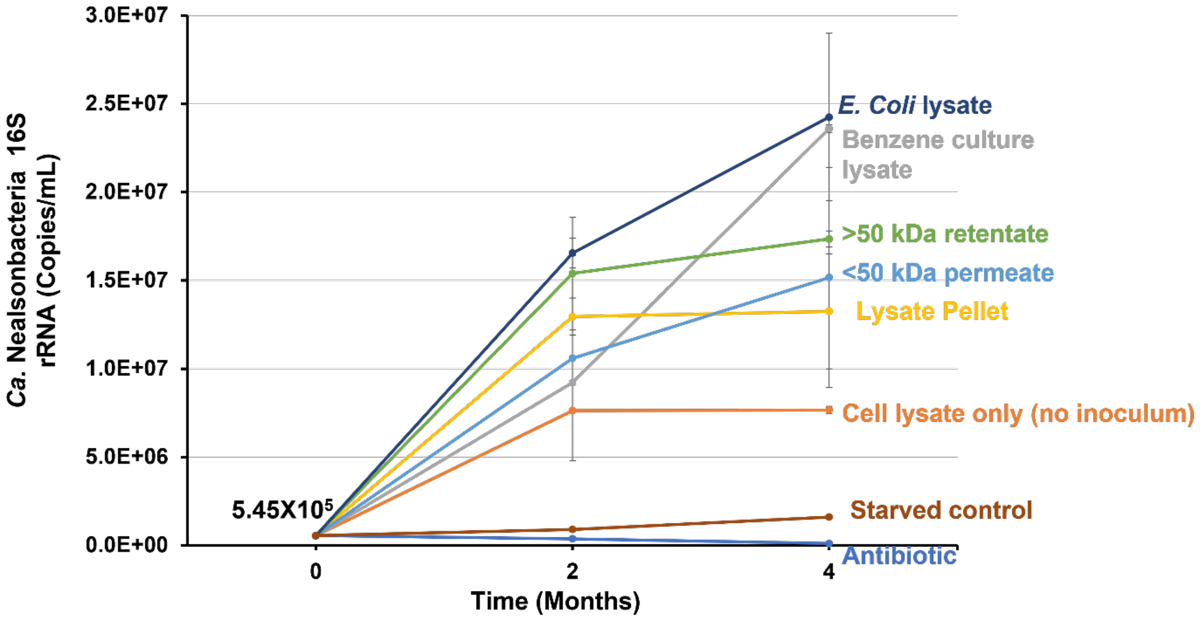


**Figure S4.** ***Ca.*** **Nealsonbacteria growth in Donor Trial #2.** One DNA sample was collected on day 1 to represent initial conditions. All other points are average of biological duplicates with error bars showing range.

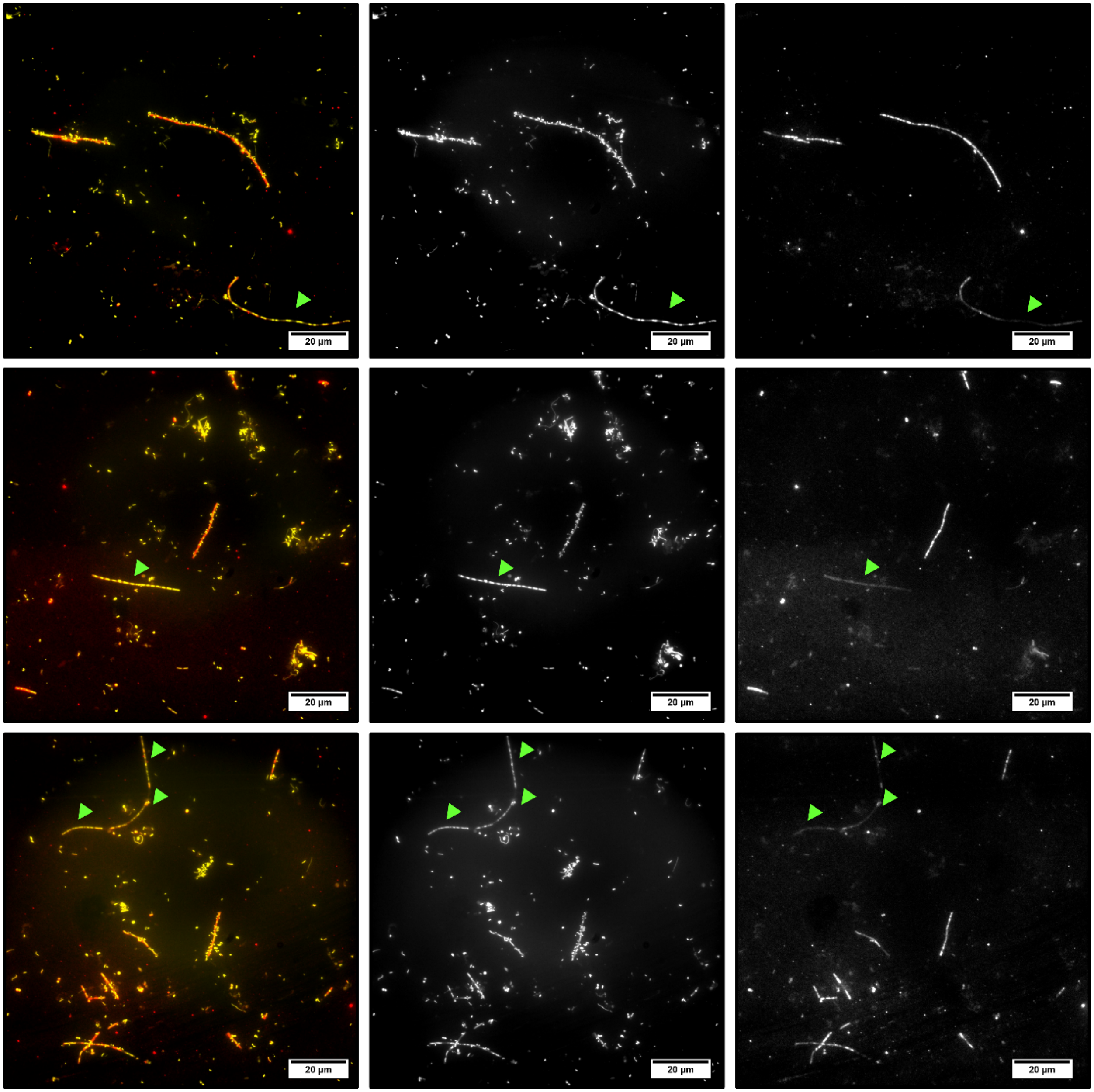


**Figure S5: Additional FISH images of *Ca.* Nealsonbacteria enrichment cultures.** The left column shows overlay images of DAPI (yellow) and Archaea probe-stained cells (red). The middle column shows DAPI stained images. The right column shows Archaea probe-stained images. Green arrows indicate the naked methanogens that were not coated with *Ca.* Nealsonbacteria cells.

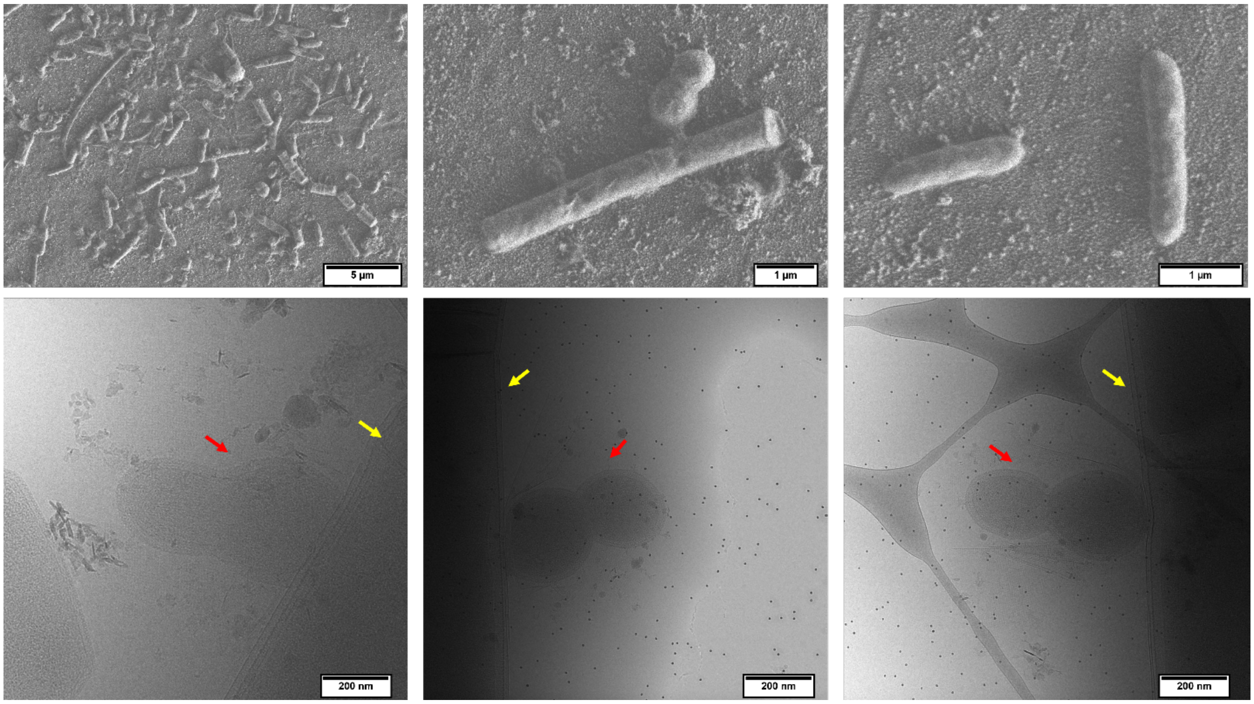


**Figure S6: Additional SEM images for methanogenic benzene-degrading culture (top row) and Cryo-EM images for *Ca.* Nealsonbacteria enrichment cultures (bottom row).** Top row shows a representative view of methanogenic benzene-degrading culture (left), *Ca.* Nealsonbacteria associated with *Methanothrix* (middle) and benzene-degrading ORM2 (right). The bottom row shows representative Cryo-EM images of *Ca.* Nealsonbacteria (red arrows) associated with *Methanothrix* (yellow arrows).

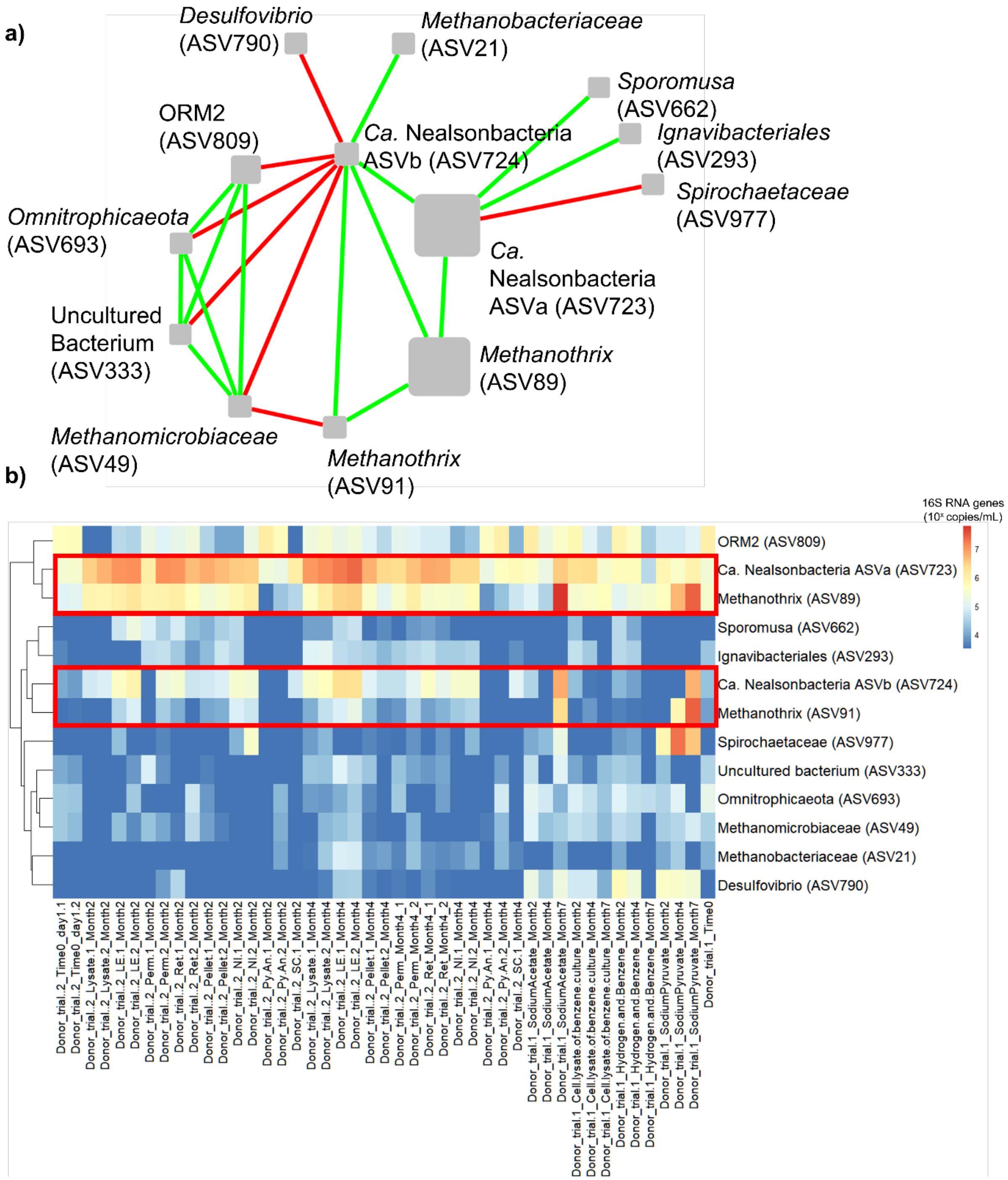


**Figure S7. Network interactions between Ca. Nealsonbacteria ASVa and ASVb and their correlated microbes (a) and corresponding heatmap (b) based on absolute abundance.** The ASV numbers in the brackets represent ordinal ASV numbers (see Table S2). The node size presents the sum abundance of each ASVs (larger size = higher abundance). Copresence (green) and mutual exclusion (red) are shown as the edges between the nodes. The rows of the heatmap were clustered using a hierarchical clustering algorithm “hclust” in R based on similarity of absolute abundance. The columns represent different treatment bottles and the names refer to Table S2. The microbes selected for the heatmap are based on the above co-occurrence network.

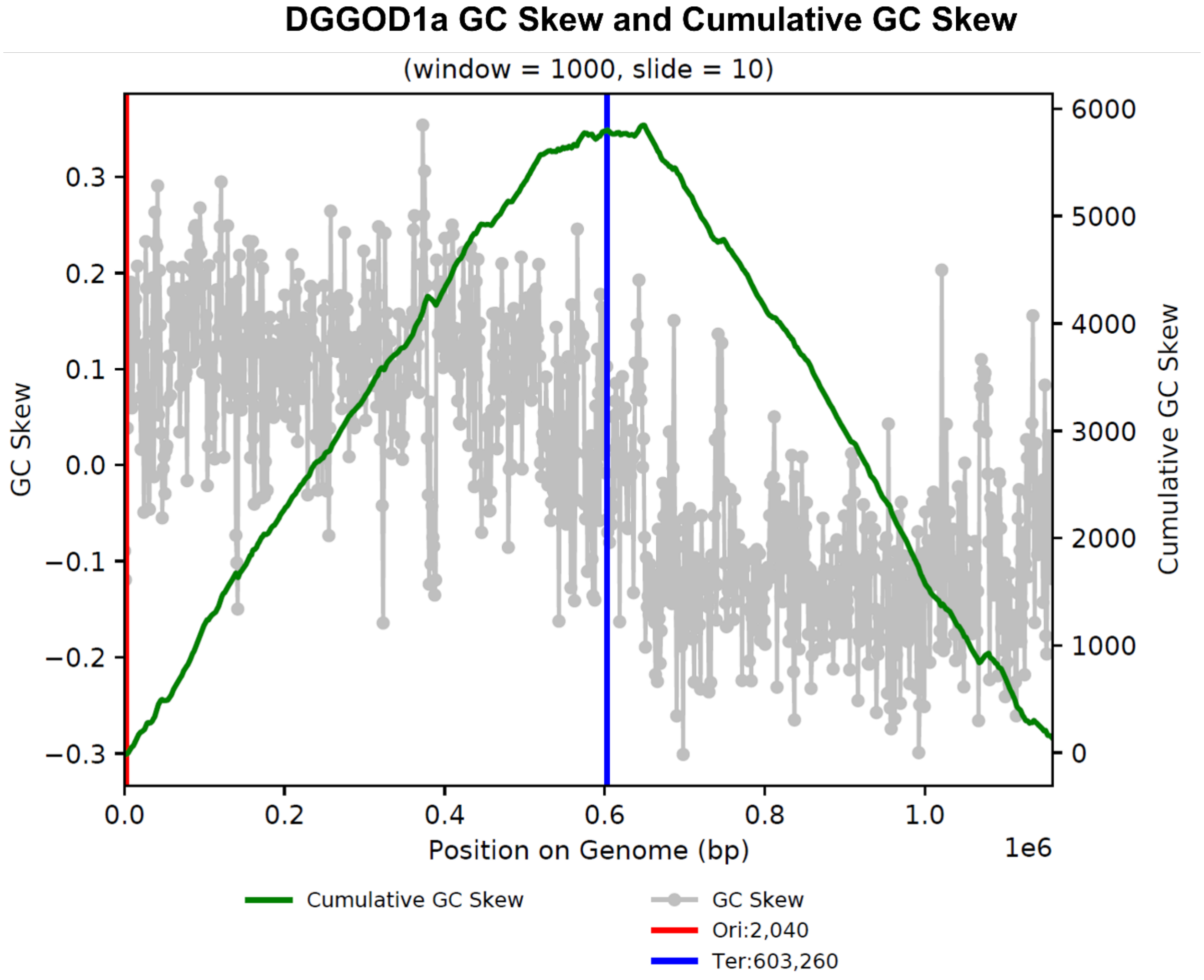


**Figure S8. GC skew and cumulative GC skew of *Ca.* Nealsonbacteria DGGOD1a genome.** The GC skew is calculated with window size of 1000 bp and slide size of 10 bp.

**Text S1: Culture lysate preparation for Donor Trials**

To prepare the lysate from the OR consortium, a 500 mL sample from OR-1bBig was concentrated ten times by tangential flow filtration using a Pellicon® XL cassette with 0.1 µm Durapore® membrane. The concentrate was then loaded into a French press cell (GlenMills 40K manual-fill cell) inside of an anaerobic glovebox (headspace containing 10% CO2, 5% H2 and 85% $N_2$), and lysed in a French press (SLM Aminco, Model FA-078) at 1500 psi. The lysate was collected in a 250 mL anaerobic glass bottle (prepared and sealed with rubber stoppers in the glove box).

For *Escherichia coli* lysate, DH5alpha cells were first grown on LB broth overnight at 37 °C. The cell pellet was collected by centrifugation at 6000 × g, and the supernatant was discarded.

To completely remove LB broth, the pellet was further washed in FeS-reduced MM medium supernatant twice, and finally resuspended in 40 mL the FeS-reduced MM medium used to grow the OR culture. The resuspended culture was placed in an anaerobic glove box for over a week to deplete any remaining oxygen by cell respiration. Cells were then lysed using a French press as above.

### Text S2: Sample pretreatment for *Ca.* Nealsonbacteria augmentation trial.

Aliquots (10 mL) from slow benzene-degrading bottle OR-p5 culture were first transferred into nine 25 ml glass vials, each amended with 15 mg/L benzene (liquid concentration) and monitored for two weeks to establish that the benzene degradation rate was consistent and low in all vials at about 0.1 mg/L/day. One of the nine vials exhibited no benzene degradation and was removed leaving 8 vials for use in the *Ca.* Nealsonbacteria augmentation experiment.

### Text S3: 16S rRNA amplicon raw read processing, NMDS and CoNet analysis.

Processing of raw 16S rRNA reads into amplicon sequence variants (ASVs) was completed using the DADA2 pipeline within sequence analysis tool QIIME2 (1, 2) and classified against the SILVA v. 132 SSU database ( https://www.arb-silva.de/documentation/release-132/) (3). Nonmetric dimensional scaling (NMDS) analysis was conducted in the R package ampvis2 (4) by using the relative abundance of all ASVs from 16S rRNA amplicon sequencing data. For the CoNet analysis, top 200 ASVs were selected and analyzed by four methods including Pearson correlation, Spearman correlation, mutual information, and Bray Curtis dissimilarity. The initial thresholds were selected so that the initial network contained 1000 positive and 1000 negative edges supported by at least three of four methods. For each method, 1000 permutation with renormalization for correlation measure and bootstrap score were computed. For bootstrapping, P-values were merged using Brown's method and the merged P-values below 0.05 were kept after using the Bonferroni correction. Edges with scores outside of the 95% confidence interval defined by bootstrap distribution or edges supported by less than three methods were discarded. The network figure was constructed with yFiles circular algorithm.

### Text S4: Metagenomic assembly, binning and functional and taxonomic annotation

Raw reads were trimmed using Trimmomatic (v 03.0) (5) with default settings to remove adaptors and to filter poor-quality reads. Assemblies were created using ABySS 1.3.7 (6) and metaSPAdes (7). The contigs were binned into metagenome assembled genomes (MAGs) using metabat2 v2.12 (8). Taxonomic classification was assigned to each MAG using the GTDB-TK tool kit which compares the genomes to the genome taxonomy database (gtdb) (9). A MAG classified as Patescibacteria according to gtdb and the superphylum Patescibacteria and phylum Parcubacteria in the NCBI taxonomy, with a size of 1,137,033 bp in 5 contigs, was selected for further refinement. Several ABySS assemblies were created using different k values to dilute out lower abundance strains. ABySS contigs were mapped onto the existing contigs and gaps in the contigs were resolved using an in-house gap resolution program described elsewhere (10, 11). The finished genome was polished by read-mapping raw reads onto the circular genome using Geneious 8.1.8 (12). Single nucleotide polymorphisms (SNPs) were edited to represent the clonal majority with 90% cut-off. The origin of replication was identified using Oriloc (13) in R. The quality of complete the genome was further evaluated by by the shape of the GC skew plot (Figure S8).

The genome was annotated using JGI/IMG (14) and a hidden Markov models search of all predicted ORFs with Pfam, TIGRfam, Panther, and custom HMM profiles from the Banfield Lab (15). The DGGOD1a genome was phylogenetically placed on a maximum likelihood concatenated ribosomal protein tree with references from across bacteria and archaea, including a large number of previously published high-quality CPR and DPANN genomes. The set of fifteen ribosomal proteins used to construct the tree (L2, L3, L4, L5, L6, L14, L15, L18, L22, L24, S3, S8, S10, S17 and S19) are encoded by a syntenic block of genes and were selected to avoid binning error chimeras. Ribosomal proteins were identified by searching predicted open reading frames against ribosomal protein databases using USEARCH. For each individual ribosomal protein, hits and reference sequences were aligned to the Pfam HMM model (16) using hmmalign from HMMer, alignments were converted from the Stockholm format to FASTA, and insertions added by hmmalign were stripped. All individual ribosomal protein alignments were concatenated together, and concatenated sequences with an ungapped length of greater than 1,100 amino acid residues were combined with reference sequences to generate the final alignment. Placement was performed with pplacer using options "-m LG -p --timing" and guppy to generate a phyloXML tree. The tree was pruned to only contain the genomes most closely related to the *Ca.* Nealsonbacteria and visualized in Mega v. 11: The Molecular Evolutionary Genetic Analysis (17). The Cluster regularly interspaced short palindromic repeat (CRISPR) loci and associated Cas proteins were identified by CRISPRCasMeta(18). The possible genomic islands were predicted by IslandViewer 4 (19).

**Text S5: Epifluorescence microscopy**

Culture samples (10 μL) were pipetted onto PTFE printed glass slides (catalog no. 63429-04 from Electron Microscopy Sciences) with wells (4 mm in diameter) for DAPI or FISH staining. Slides were prepared using a standard FISH protocol (20-24). The details of hybridization conditions are described in Text S6. Slides were covered with glass coverslips and sealed with nail polish. A droplet of immersion oil was added to each slide and the slides were observed using an epifluorescent microscope (BX 51, Olympus) with a 100 × UPlan Apochromat objective, 150 W xenon lamp (Opti Quip), a set of excitation filter cubes (DAPI/Cy3/Cy5) and 10× focusing eyepiece to check for uniformity of cell morphology (1,000-fold magnification). The microscopic images were captured by Hamamatsu ORCA-Flash 4.0 digital CMOS camera. All of the *Ca.* Nealsonbacteria specific probes designed and tested in this study are shown in Table S4, however, none of them showed a positive binding signal other than background noise.

**Text S6: Fluorescence in situ hybridization (FISH) method detail**

FISH probes used in this study are shown in Table S4 (Excel). Although these probes eventually were found not to work, the details of these trials are provided for anyone to review and improve. The design of FISH probes for *Ca.* Nealsonbacteria was conducted using the phylogeny software ARB (25). The 16S rRNA gene sequence of *Ca.* Nealsonbacteria DGGOD1a was manually added into the SILVA non-redundant 16S rRNA database. All potential probes with GC content over 50% were selected and BLAST searched on NCBI (Jan. 2018) to check for specificity. All the probes which passed this screening were analyzed using the oligonucleotide probe evaluation tool mathFISH (26, 27) to find an optimal formamide concentration.

All samples for FISH were first harvested by centrifuge of an aliquot of 2 ml culture at 13000 × g for 20 min. The supernatant was then removed, and the pellet was resuspended in 950 µL 1× phosphate buffered saline (PBS). A 50 µL of 20% paraformaldehyde (PFA) fixative was then added to a 1% final concentration. The mixture was then left in a 4°C fridge for at least 24 hours incubation. After incubation, the mixture was pelleted down at 13000 × g for 20 min and resuspended with 500 µL PBS, twice, to wash off the fixative. An aliquot of 10 µL of the fixed cell was then loaded onto a PTFE printed glass with wells in 4 mm diameter, and let it air-dry. The air-dried glass slides were then dehydrated for 3 min each in 50, 80, and 100% ethanol.

The next steps are hybridization with the probe, and then washing (20-24). Details of these two critical steps are outlined below.

The hybridization mixture was prepared by mixing 1 volume of probe solution (50 ng/µL) to 9 volumes of hybridization buffer in a 0.5 ml microfuge tube. The detailed hybridization buffer compositions are shown in Table A (and subsequent associated Tables B-D). A 50 mL polyethylene centrifuge tube was used as the hybridization vessel. A piece of Kimwipes paper was soaked with hybridization buffer and inserted into the centrifuge tubes. Separate tubes were used for different formamide concentrations.

**Table A. Hybridization buffer compositions**

| Reagent | Reagent stock solution concentration | Volume (µL) | Final concentration |
|---:|---:|---:|---:|
| NaCl | 5 M | 360 | 900 mM |
| Tris/HCl | 1 M | 40 | 20 mM |
| SDS | 20% (w/v) | 1 | 0.01% (w/v) |
| Formamide | Pure | Depended on probe used (see Table S4 and Table B below) | |
| milliQ $H_2O$ | | Added to 2 mL | |

**Table B. Formamide and water reference table**

| Formamide concentration | Formamide volume (µL) | milliQ water volume (µL) |
|---|---|---|
| 0% | 0 | 1598 |
| 5% | 100 | 1498 |
| 10% | 200 | 1398 |
| 15% | 300 | 1298 |
| 20% | 400 | 1198 |
| 25% | 500 | 1098 |
| 30% | 600 | 998 |
| 35% | 700 | 898 |
| 40% | 800 | 798 |
| 45% | 900 | 698 |
| 50% | 1000 | 598 |
| 55% | 1100 | 498 |
| 60% | 1200 | 398 |

Ten (10) µL of hybridization mixture was loaded into the well to fully cover the sample, and the slides were inserted into the centrifuge tubes in a horizontal position. The centrifuge tubes were then incubated in a 46°C incubator for two hours. Meanwhile, the washing buffer was prepared in a 50 mL polyethylene centrifuge (details of composition in Table C) and preheated in a 48 °C water bath.

**Table C. Washing buffer composition**

| Reagent | Reagent stock solution concentration | Volume | Final concentration |
|---|---|---|---|
| NaCl | 5 M | Depended on % formamide in hybridization buffer (see Table A and Table D) | Depended on % formamide in hybridization buffer (see Table A and Table D) |
| Tris/HCl | 1 M | 1 mL | 20 mM |
| SDS* | 20% (w/v) | 25 µL | 0.01% |
| EDTA** | 0.5 M | 500 µL | 5 mM |
| milliQ $H_2O$ | Pure | Added to 50 mL | |

*Added SDS at the last step to avoid precipitation

**Only if 20% formamide or more was used

**Table D. NaCl concentration reference table (for 50 mL of washing buffer)**

| Formamide concentration in hybridization buffer | NaCl Concentration (M) | 5M NaCl volume (µL) |
|---|---|---|
| 0 | 0.900 | 8900 |
| 5 | 0.636 | 6260 |
| 10 | 0.450 | 4400 |
| 15 | 0.318 | 3080 |
| 20 | 0.225 | 2150 |
| 25 | 0.159 | 1490 |
| 30 | 0.112 | 1020 |
| 35 | 0.080 | 700 |
| 40 | 0.056 | 460 |
| 45 | 0.040 | 300 |
| 50 | 0.028 | 180 |
| 55 | 0.020 | 100 |
| 60 | 0.014 | 40 |
| 65 | - | - |
| 70 | - | - |

*Addition of EDTA contributes to the $Na^+$ concentration, therefore, the required volume of 5 M NaCl solution in the washing buffer was reduced by 100 uL.

After two hours of incubation, the slides were rinsed with washing buffer and transferred into preheated washing buffer, and incubated for 25 min in a 48°C water bath. After the water bath, the slides were rinsed with sterile Milli-Q water and air-dried. Then, the air-dried slides were stained with 10 µL of a 1 µg/mL DAPI solution each well for 3 min followed by rinsing and air drying.

The stained slides were then mounted in a 4:1 mixture of Citifluor (Citifluor Ltd, London, UK) and Vecta Shield (Vector Laboratories, Inc., Burlingame, CA), covered with glass coverslips and sealed with nail polish.

**Text S7: Ionic liquid sample preparation and SEM setup**

Concentrated biological samples (microbial cell pellets from centrifugation at 13,000 × g for 20 min) were first fixed in 2% glutaraldehyde in cacodylate buffer and rinsed with Milli-Q water. After fixation, the pellet was re-suspended in a 5% Hitachi HiLEM IL1000 ionic liquid (IL) $C_7H_{19}NO_4S$ (28) water solution at room temperature. The mixture was then passed through a 0.02 µm cut-off membrane (catalog no. CA28138-061, WhatmanTM Anopre Inorganic Membrane). After filtration, the membrane was placed on an aluminum sample stub for 12 hours to ensure sufficient equilibrium of IL and intracellular liquid. Excess IL was carefully rinsed off with water and the membrane was placed on the stub and observed by SEM. SEM was carried out using a Hitachi SU8230 cold FEG SEM at 0.5 and 1 kV at Ontario Centre for Characterization of Advanced Materials (OCCAM) of the University of Toronto.

**Text S8: Cryo-EM sample preparation and operational condition**

Small (2 mL) samples from the parent culture of Donor Trial #1 and the bottle amended with cell lysate of benzene culture after four months since inoculum in Donor Trial #1 were collected in screw-top microcentrifuge tubes with o-rings. Samples were concentrated four times by centrifugation at 13,000 × g for 20 min. All sample handling was conducted under anaerobic conditions. The samples were then shipped to UC Berkeley for Cryo-EM analysis. The concentrate was deposited onto lacey carbon-coated formvar Cu-grids (200 mesh and 100 microns, Electron Microscopy Sciences LC200-CU-100) and plunge frozen into liquid propane at liquid nitrogen temperature using a Vitrobot Mark IV (blot force two for one second). Colloidal gold particles (250 nm) were put on grids before sample addition. Grids were then stored in liquid nitrogen until analysis. Cryo-EM images were captured on a 300 kV JEOL3100 with in column energy filter with a Gatan K2 DED.

**Tables S1-S5, S8-S10** are larger and are provided in the associated Excel file.

**Table S6: Specific FISH probes used/tested in this study. (NB: None of the OD1 Probes worked)**

| **Probe** | **Specificity** | **Probe Sequence (5' - 3')** | **Formamide concentration (%)*** | **5'-Modification** | **Purification** | **Reference** |
|---|---|---|---|---|---|---|
| Arch915 | Archaea | GTGCTCCCCCGCCAATTCCT | 35 | Cyanine 5 | HPLC | (35) |
| OD-81 | *Ca.* Nealsonbacteria in this study | ACCCTGCACGGGAAAATC | 0 | Cyanine 3 | HPLC | This Study |
| OD-412 | *Ca.* Nealsonbacteria in this study | ACGGCGCGAACGTCTTCA | 30 | Cyanine 3 | HPLC | This Study |
| OD-501 | *Ca.* Nealsonbacteria in this study | GCACAGAATTAGCCCCCG | 20 | Cyanine 3 | HPLC | This Study |
| OD-622 | *Ca.* Nealsonbacteria in this study | CCGGTTCACCCCCCGGGT | 20 | Cyanine 3 | HPLC | This Study |
| OD-736 | *Ca.* Nealsonbacteria in this study | TCAAGATTGCCCCAGCTA | 20 | Cyanine 3 | HPLC | This Study |
| OD-996 | *Ca.* Nealsonbacteria in this study | CGGGACGATCAGTAGAAT | 0 | Cyanine 3 | HPLC | This Study |
| OD-1209 | *Ca.* Nealsonbacteria in this study | CGGCCCCAGTTGTCAAAG | 20 | Cyanine 3 | HPLC | This Study |

*Formamide concentration was estimated using Math-FISH formamide curves at 46°C, 0.9M [Na+] and probe concentration 250 nM, ± 5% wash test for each probe to find the optimal formamide concentration.

**Table S11: Specific 16S rRNA primers used for qPCR in this study**

| Primer Name | Target Organism | Primer Sequence (5' - 3') | Expected Amplicon Length | Primer Set Specificity* | Annealing Temperature (°C) | References |
|---|---|---|---|---|---|---|
| Bac_1055f | General Bacteria | ATGGCTGTCGTCAGCT | 338 | 1085282/3195888 (kingdom Bacteria) | 55 | (29, 30) |
| Bac_1392r | | ACGGGCGGTGTGTAC | | | | |
| Arch_787f | General Archaea | ATTAGATACCCGBGTAGTCC | 273 | 56410/160768 (kingdom Archaea) | 59 | (31) |
| Arch_1059r | | GCCATGCACCWCCTCT | | | | |
| ORM2_168f | Thermodesulfobacteria, ORM2 | GAGGGAATAGCCAAAGGTGA | 274 | 15/3195888 (kingdom Bacteria) | 59 | (32, 33) |
| ORM2_422r | | GAGCTTTACGACCCGAAGAC | | 16/14443 (unclassified Deltaproteobacteria spp.) | | |
| OD1_987f | The dominant *Ca.* Nealsonbacteria in the OR/DGG consortium | GGTGCTGCATGGTTGTCGTC | 200 | 47501/3195888 (kingdom Bacteria) | 65 | (34) |
| OD1_1186r | | GCTGCCCTCTGTAACTGCCA | | | | |

*The 16S rRNA primer specificity was determined based on Ribosomal Database Project (RDP) Probe Match tool release 11, Update 5 with a maximum of two nucleotide differences allowed.

**Table S12: Summary of all qPCR standard curve equations and efficiencies in this study used to quantify total Bacteria, total Archaea, Thermodesulfobacteria ORM2, and *Ca.* Nealsonbacteria DGGOD1a.**

| **Total Bacteria** | | | |
|---|---|---|---|
| Efficiency (%) | $R^2$ | Slope | Y-int |
| 100.8 | 0.999 | -3.302 | 33.657 |
| 96.7 | 0.999 | -3.403 | 33.917 |
| 96.8 | 0.999 | -3.402 | 34.378 |
| 103.8 | 0.997 | -3.235 | 33.485 |
| 99.1 | 0.998 | -3.343 | 33.898 |
| 98.2 | 0.998 | -3.367 | 34.302 |
| 94.7 | 0.998 | -3.455 | 34.313 |
| **Total Archaea** | | | |
| Efficiency (%) | $R^2$ | Slope | Y-int |
| 92.9 | 0.999 | -3.504 | 35.106 |
| 96.9 | 0.999 | -3.408 | 36.133 |
| 92.8 | 0.995 | -3.507 | 36.876 |
| **Thermodesulfobacteria ORM2** | | | |
| Efficiency (%) | $R^2$ | Slope | Y-int |
| 92.9 | 0.998 | -3.505 | 35.119 |
| 97.7 | 0.998 | -3.387 | 34.27 |
| 89.7 | 0.998 | -3.597 | 36.495 |
| 95.0 | 0.999 | -3.447 | 35.788 |
| 96.0 | 0.998 | -3.422 | 34.546 |
| 100.0 | 0.998 | -3.323 | 34.239 |
| 98.2 | 0.991 | -3.366 | 34.377 |
| ***Ca.* Nealsonbacteria DGGOD1a** | | | |
| Efficiency (%) | $R^2$ | Slope | Y-int |
| 92.4 | 0.998 | -3.519 | 34.611 |
| 94.1 | 0.999 | -3.471 | 35.683 |
| 95.6 | 0.999 | -3.432 | 35.818 |
| 91.7 | 0.999 | -3.537 | 36.172 |
| 91.9 | 0.998 | -3.532 | 34.802 |
| 99.9 | 0.998 | -3.325 | 35.671 |
| 102.9 | 0.999 | -3.255 | 34.772 |
| 94.4 | 0.996 | -3.463 | 36.393 |
| 98.3 | 0.999 | -3.364 | 35.487 |

14. Grigoriev IV, Nordberg H, Shabalov I, Aerts A, Cantor M, Goodstein D, Kuo A, Minovitsky S, Nikitin R, Ohm RA, Otillar R, Poliakov A, Ratnere I, Riley R, Smirnova T, Rokhsar D, Dubchak

I. 2012. The genome portal of the Department of Energy Joint Genome Institute. Nucleic Acids Res 40:D26-32.
